# Oral Exposure to Benzalkonium Chlorides in Male and Female Mice Reveals Sex-Dependent Alteration of the Gut Microbiome and Bile Acid Profile

**DOI:** 10.1101/2024.05.13.593991

**Authors:** Vanessa A. Lopez, Joe L. Lim, Ryan P. Seguin, Joseph L. Dempsey, Gabrielle Kunzman, Julia Y. Cui, Libin Xu

## Abstract

Benzalkonium chlorides (BACs) are commonly used disinfectants in a variety of consumer and food-processing settings, and the COVID-19 pandemic has led to increased usage of BACs. The prevalence of BACs raises the concern that BAC exposure could disrupt the gastrointestinal microbiota, thus interfering with the beneficial functions of the microbes. We hypothesize that BAC exposure can alter the gut microbiome diversity and composition, which will disrupt bile acid homeostasis along the gut-liver axis. In this study, male and female mice were exposed orally to d_7_-C12- and d_7_-C16-BACs at 120 µg/g/day for one week. UPLC-MS/MS analysis of liver, blood, and fecal samples of BAC-treated mice demonstrated the absorption and metabolism of BACs. Both parent BACs and their metabolites were detected in all exposed samples. Additionally, 16S rRNA sequencing was carried out on the bacterial DNA isolated from the cecum intestinal content. For female mice, and to a lesser extent in males, we found that treatment with either d_7_-C12- or d_7_-C16-BAC led to decreased alpha diversity and differential composition of gut bacteria with notably decreased actinobacteria phylum. Lastly, through a targeted bile acid quantitation analysis, we observed decreases in secondary bile acids in BAC-treated mice, which was more pronounced in the female mice. This finding is supported by decreases in bacteria known to metabolize primary bile acids into secondary bile acids, such as the families of Ruminococcaceae and Lachnospiraceae. Together, these data signify the potential impact of BACs on human health through disturbance of the gut microbiome and gut-liver interactions.

## Introduction

Benzalkonium chlorides (BACs) are a subclass of quaternary ammonium compounds (QACs) that are widely used because of their broad-spectrum antimicrobial properties (Arnold *et al*., 2023a). BACs are used in a range of applications from residential, agricultural, industrial (food processing equipment and milking equipment), and clinical (hand sanitizers, eye, and nasal drops) settings (Pereira and Tagkopoulos, 2019). High levels of BACs have been reported in several food products, including grapefruit seed extracts, fruits and vegetables, and dairy products. Notably, the median and maximum levels of BACs in milk were 0.15 and 6.7 mg/kg, respectively, with even higher median and maximum levels measured in milk-based ice cream at 0.22 and 22 mg/kg, respectively (Takeoka *et al*., 2005; The Federal Institute for Risk Assessment (BfR) of Germany, 2012a; Slimani *et al*., 2017); BACs were also detected as an unpermitted ingredient in food additives applied in meat products (Kröckel *et al*., 2003). Additionally, direct application of BACs is performed on eating utensils, egg shells, milking equipment and udders, and medical instruments (US EPA, 2006a). Given the ubiquitous environmental presence of BACs, humans may be chronically and systemically exposed by several routes: direct dermal/eye contact, inhalation, and ingestion (US EPA, 2006).

Literature suggests that BACs can be absorbed into the body, and exposure amongst human populations is significant. For example, BACs are readily detectable in human blood following acute oral ingestion (Xue *et al*., 2002; Mishima-Kimura *et al*., 2018). In a fatal ingestion case of an 83-year-old man, BACs were measured at 245-264 nM in the blood 18 days post-ingestion, thereby suggesting persistence in the body (Mishima-Kimura *et al*., 2018). We reported BAC detection in the blood of 80% of human participants—half of whom had BAC concentrations in the 10-150 nM range (Hrubec *et al*., 2021). Due to the COVID-19 pandemic, excessive usage of BAC-containing disinfectants like Clorox and Lysol products has increased environmental exposure to BACs well above pre-pandemic levels. For example, the quantitation of QACs in indoor residential dust suggests an almost doubling of the levels of BACs during COVID-19 relative to pre-COVID-19 (Zheng *et al*., 2020). Additionally, Zheng et al. showed median levels of BACs in human blood increased 2.7-fold during the COVID-19 pandemic relative to pre-COVID-19 (Zheng *et al*., 2021). The prevalent use of BACs as antimicrobials raises the concern of potential disruption to gut microbiota homeostasis. Previous studies in rat models have shown fecal excretion as a major route of elimination of the BACs even when administered through IV injection (Luz *et al*., 2020). More recently, we detected BACs and BAC metabolites in five human fecal samples, reaching over 1 μM for total BAC concentrations (Nguyen *et al*., 2024). Thus, the interaction of BACs with gut microbiota is highly likely regardless of the exposure route.

The gastrointestinal microbiome has become increasingly recognized as an invaluable contributor to human (host) health. The gut microbiome is defined as a diverse consortium of commensal, symbiotic, and pathogenic microorganisms. These organisms include bacteria, archaea, fungi, protozoa, and viruses that inhabit the gut (Barko *et al*., 2018), with bacteria being the main organism (Sirisinha, 2016). The symbiotic relationship between the host and the gut microbiome has been linked with beneficial contributions to overall host health, like immunity, nutrition, and human development (Eloe-Fadrosh and Rasko, 2013), while gut microbiome dysbiosis was found to contribute to many human diseases, such as obesity, asthma, some cancers, heart disease, and autism (Barko *et al*., 2018).

The gut microbiome establishes its relationship with the host by participating in various axes. Crosstalk between the gut and liver has become increasingly recognized by parallel rises in occurrences of liver diseases and GI and immune disorders (Tripathi *et al*., 2018). The gut and liver communicate via tight bidirectional links through the biliary tract, portal vein, and systemic circulation. Bile Acids (BAs) are a group of steroidal acids derived from cholesterol in hepatocytes and are imperative signaling molecules for intermediary metabolism within the gut-liver axis (Li *et al*., 2018). Furthermore, dysbiosis in the gut microbiome shifts the balance between primary and secondary BAs and their subsequent enterohepatic recycling (Figure 1).

**Figure 1.**
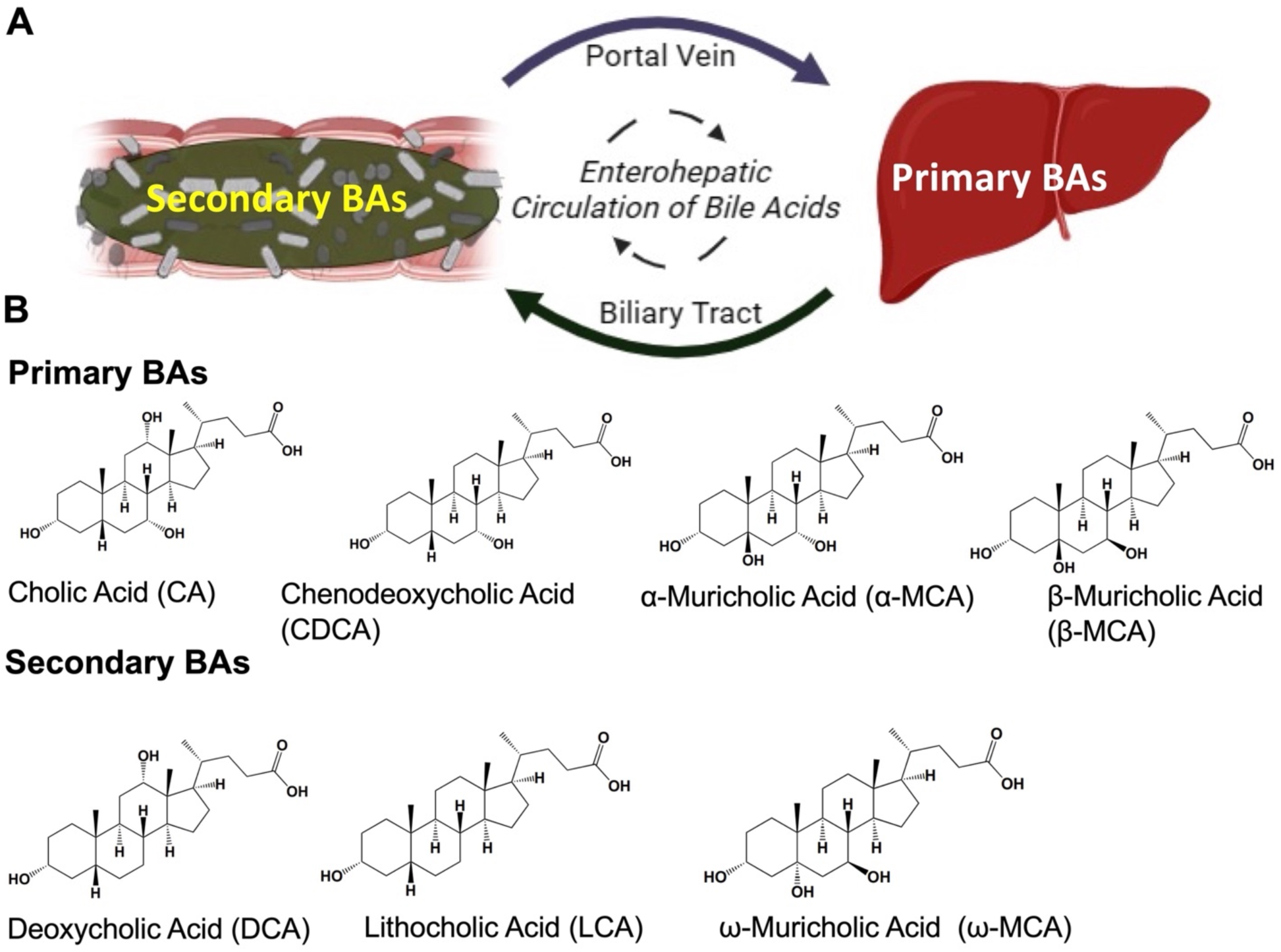
Schematic of Enterohepatic Circulation of Bile Acids. **A.** Primary BAs, cholic acid (CA) and chenodeoxycholic acid (CDCA) are formed from cholesterol in hepatocytes (Hofmann, 1984). The muricholic acid (MCA) family belongs to rodents (Hofmann, 2009,) of which α- and β-MCA are primary bile acids. The primary BAs become conjugated to taurine or glycine to increase their hydrophilicity and enter the biliary tract (Staels and Fonseca, 2009). After brief storage in the gallbladder, the conjugated primary BAs travel through the biliary tract to the small intestine and eventually reach the distal ileum. Conjugated BAs can get absorbed by the portal vein and travel back to the liver for secretion to the bile (Stellaard and Lütjohann, 2021). The unabsorbed BAs will undergo modification by bacteria, such as a) deconjugation; b) dehydrogenation and c) dehydroxylation (Hofmann, 2009), forming “secondary” BAs. The following secondary BAs, lithocholic acid (LCA), deoxycholic acid (DCA) and ω-muricholic acid (ω-MCA), are formed from the respective primary BAs. BAs enter the portal vein by either passive diffusion in the small intestine and colon or active transport in the distal ileum (Winston and Theriot, 2020), followed by uptake by hepatocytes. **B.** Chemical structures of BAs quantified in this study; from left to right; first panel CA, CDCA, α-MCA, β-MCA; second panel DCA, LCA, ω-MCA.

An imbalance in BAs and gut bacteria can elicit a cascade of host immune responses relevant to the progression of liver diseases (Tripathi *et al*., 2018). Previous work (L. Zhang *et al*., 2015a; S. Zhang *et al*., 2015a; Fazeli *et al*., 2011a; Lu *et al*., 2014a; Breton *et al*., 2013a; Claus *et al*., 2016a; Naspolini *et al*., 2022a) has elucidated the potential of environmental toxicant exposure on microbiome composition. The objective of this study was to assess the hypothesis that BAC exposure could alter gut microbiome composition and diversity and subsequently affect bile acid homeostasis between the gut and liver.

## Materials and Methods

### Animals

Seven to eight-week-old C57BL/6J male and female mice were purchased from Jackson Laboratories (Bar Harbor, Maine). Experiments were staggered, with male mice undergoing the exposure protocol first. The University of Washington Institutional Animal Care and Use Committee approved all animal protocols. All experiments followed the Guiding Principles in the Use of Animals in Toxicology. Mice were acclimated to the animal facility for 2 weeks. Mice were then acclimated to an untreated Nutra-Gel diet (Product F5769-KIT, Bio-Serv, Frenchtown, New Jersey) for 2 weeks before BAC exposure. Deuterated BACs were used to ensure prevention of interruption of accurate quantitation, as described in Herron et al., 2018. Mice were randomly assigned to exposure groups (n = 6): control Nutra-Gel diet, or treatment with d_7_-C12-BAC (120 µg/g/day) or d_7_-C16-BAC (120 µg/g/day) for 1 week. At the end of the first week, mice were sacrificed, and the following tissues and fluids were collected, flash-frozen in 2-methylbutane on dry ice and stored at -80 °C until subsequent analyses: liver, ileum, jejunum, duodenum, including intestinal content in all three intestinal tissue section, and large intestine (colon and cecum), and blood via cardiac puncture. Feces were collected throughout the Nutra-gel acclimation period, as well as the treatment period (days 1, 6, 15, 18, and 21). Feces were kept at -80°C until subsequent analysis.

### Chemicals

Optima LC/MS solvents (acetonitrile, chloroform, methanol, and water), 2-methylbutane, and formic acid were purchased from Thermo Fisher Scientific (Grand Island, New York). d_7_-C12-BAC and d_7_-C16-BAC were prepared as described previously (Herron et al., 2016). The deuterated (d_4_-) bile acid standards, including tauro-α-muricholic acid-d_4_ (sodium salt) (T-α-MCA), α-muricholic acid-d_4_ (α-MCA), and ω-muricholic acid-d_4_ (ω-MCA), were purchased from Cayman Chemical (Ann Arbor, Michigan). Bile acids, including cholic acid (CA), chenodeoxycholic acid (CDCA), deoxycholic acid (DCA), lithocholic acid (LCA), α-muricholic acid (α-MCA), β-muricholic acid (β-MCA), and ω-muricholic acid (ω-MCA), were purchased from Cayman Chemical (Ann Arbor, Michigan).

### Bacterial DNA Isolation and 16S rRNA Sequencing

Total DNA was extracted from the cecum of untreated and treated mice using E.Z.N.A. DNA Stool Kit (Omega Bio-tek, Inc., Norcross, GA) according to the manufacturer’s protocol. The concentration of DNA was determined via Qubit 2.0 Fluorometer (Life Technologies/Thermo Fisher Scientific, Grand Island, NY). The integrity and quality of DNA samples were confirmed by the Agilent 2100 Bioanalyzer (Agilent Technologies Inc., Santa Clara, CA). The V4 region of the 16S rRNA gene was amplified and sequenced using HiSeq. 2500 platform (250-bp paired-end) (Novogene, Beijing, China) (n = 4). The paired-end sequence reads were merged, demultiplexed, and filtered using QIIME2 version 2020. 2 (Caporaso et al., 2010). Alpha diversity was assessed by QIIME2-2020.2. at the p-max depth of 10,000. The input table was acquired from qiime dada2 denoise paired steps. Input phylogeny came from the acquisition of a rooted phylogenetic tree using QIIME2-2020.2. Beta diversity was examined by a weighted unifrac distance matrix from core metric results acquired. Taxa composition (bacterial phyla to species level) was defined by Silva132. Taxa data acquired by QIIME2-2020.2 were used for analysis at different levels (phylum, family, and genera; see Supplemental Excel file).

### Bile Acid and BAC Extractions

A standard stock solution of each BA and deuterated IS was prepared at a concentration of 1 mg/ml in methanol. A 5% NH_4_OH in Acetonitrile solution containing each d_4_-BA at a concentration of 0.175 µg/mL was spiked into each sample for LC/MS-MS analysis.dd

For fecal BA extraction, feces were accurately weighed and stored at -80 °C until the time of BA extraction. 2.5 µL of MilliQ water was added for every mg of feces. Feces-water mixtures were sonicated in ice water for 30 minutes to produce feces-water homogenates. 10 µL of ice-cold alkaline acetonitrile (ACN) (5% ammonia in ACN) with IS mixture (d_0_-C12 and C16 BACs, ω-OH C12- and C16-BAC, COOH metabolites 1µM each; d_4_-bile acids 0.175 µg/mL) for every mg of feces was added to the homogenate, and vortexed vigorously for 5-8 seconds and left to equilibrate on ice for 10 minutes. Samples were sonicated for 1 hour and then centrifuged at 12, 000 x g for 15 min at 4 °C. Supernatants were collected in new tubes, and pellets were resuspended in 750 µL of 100% methanol. Samples were sonicated for 20 minutes and then centrifuged at 12,000 x g for 20 minutes to isolate a second supernatant. Two supernatants were combined and evaporated by SpeedVac Vacuum (30 °C) for 3-5 hours. Samples were reconstituted in 100 µL of 1:1 methanol:water mix. The suspension was transferred into 0.2 µm costar Spin-X HPLC microcentrifuge filter tubes and centrifuged at 12, 000 x g for 10 minutes. Filtrate (70 µL) was transferred to LC–MS vials and stored for analysis.

For liver BA extraction, 60-70 mg of liver were accurately weighed. Frozen liver tissues were homogenized in 5 volumes of MilliQ water by Bead Homogenizer. Homogenates were transferred from homogenization tubes to clean tubes. 10 µl of cold alkaline Acetonitrile (ACN) (5% ammonia in ACN) with IS mixture (d_0_-C12- and C16-BACs, ω-OH C12- and C16-BAC, COOH metabolites 1µM each; d_4_ bile acids 0.175 µg/mL) for every mg of liver tissue weight was added. This extraction process extracted bile acids, as well as BAC and BAC metabolites. Tubes were vortexed and then sonicated for 1 hour at room temperature. Samples were centrifuged at 12,000 g for 15 minutes at 4℃. Supernatants were transferred to clean tubes, and 750 µL MeOH was added to the pellets. The mixture was sonicated for 30 minutes and then centrifuged at 15,000 g for 20 minutes. Supernatants were combined and dried under SpeedVac. The residue was reconstituted in 100 µL of 50% MeOH, and samples were vortexed and transferred into a 0.2 µm Costar Spin-X centrifuge tube filter. Samples were centrifuged at 12,000g for 10 minutes. Filtrate (70 µL) was transferred to LC-MS vials and stored for analysis.

For male and female blood BAC extractions, 10 μL of thawed whole blood was added to 20 μL of water and sonicated for 15-20 minutes. 10 μL of ACN spiked with d_0_-BAC Internal Standard series (C6, C8, C10, C12, C14 and C16 d_0_-BACs; C8, C10, C12, C14 and C16 d_0_-COOH-BACs; d_0_-ω-OH-C12 and d_0_-ω-OH-C16 at 40 nM each). Samples were vortexed and left to sit on ice for 5-10 minutes. 100 µL of ACN: MeOH (1:1) was added to each sample. Samples were vortexed and left to rest on ice for 5-10 minutes. Samples were centrifuged at 4°C for 15 minutes, and sample supernatants were transferred to LC-MS vials and stored for analysis.

### Targeted Analysis of BAs, BAC, and BAC metabolites

Bile acids were analyzed by ultra-high-performance liquid chromatography-tandem mass spectrometry (UHPLC-MS/MS) using a triple-quadrupole mass spectrometer (Triple-Quad 6500+; SCIEX, Vaughan, Ontario, Canada) equipped with electrospray ionization (ESI) by modifying previous methods (Y. Zhang and Klaassen, 2010; Qiu *et al*., 2016). 5 μl of each prepared sample was injected into the system. Reverse chromatographic separation was achieved on a Thermo Hypersil GOLD C18 column (100 x 2.1 mm, 1.9 µm particle size) under the following conditions; flow rate, 0.400 ml/min, and gradient elution method with solvent A (20% acetonitrile/80% H_2_O with 0.1% formic acid) and solvent B (20% H_2_O/80% acetonitrile with 0.1% formic acid), 0 min, 5% B; 2 min, 5% B; 4 min, 8% B; 8 min, 30% B; 12 min, 80% B; 13.5 min, 90% B; 15 min, 80% B; 16 min, 5% B; 19 min, 5% B. (MS conditions shown in Supplemental Table 1).

Selective reaction monitoring (SRM) was used to monitor mass-to-charge ratios (m/z) of the parent ion (Q1) for each respective bile acid and their respective characteristic fragment (Q3). Mass transitions are shown in Supplemental Table 2. Utilization of both positive and negative modes was performed for this assay. Analyst software was used to integrate extracted BA peaks. Calibration curves were constructed for each BA internal standard, and BA concentrations were calculated based on the ratio of the analyte peak area to the internal standard peak area. β-MCA was ratioed to the average of d_4_-α-MCA and d_4_-ω-MCA; d_0_-CDCA was ratioed to d_4_-CA. All data are presented as the median. Statistical analyses were performed on GraphPad Prism (GraphPad Software, La Jolla, CA) using Welch’s one-way ANOVA analysis followed by Dunnett’s multiple comparison tests relative to the Control.

BAC and BAC metabolites were analyzed by ultra-high-performance liquid chromatography-tandem mass spectrometry (UHPLC-MS/MS) using a Synapt G2-XS ion mobility Q-TOF mass spectrometer equipped with electrospray ionization (ESI) in the positive mode (Waters Corporation, Milford, MA). 5-10 µl of each prepared sample was injected into the system. Reverse chromatographic separation was achieved on a Thermo Hypersil GOLD C18 column (100 x 2.1 mm, 1.9 µm particle size) at ambient temperature, with a flow rate of 0.4 ml/min. ACUITY UPLC system and autosampler (Waters Corporation, Wilford, MA) were used for mobile phase delivery and sample injection. The solvent gradient comprised of mobile phase A: 0.1% formic acid, 2mM ammonium formate in water, and solvent B: Acetonitrile: 0 min: 15% B, 14 min: 85%, 15-17 min: 100% B, 18.5-20 min: 15%B. ESI conditions were as described in (Vieira *et al*., 2024a). Supplemental Table 8 describes all Analyte retention times and precursor to product ion *m/z* transitions. Because the C57BL/6J mice were dosed with a deuterated BAC structure (d_7_), use of the following d_0_-BACs were utilized as internal standards (COOH C_8_-BAC, COOH C_10_-BAC, COOH C_12_-BAC, COOH C_16_-BAC, C_12_-BAC and C_16_-BAC). The TargetLynx application in MassLynx (Waters Corporation) was utilized for peak integrations and analysis.

Analyte peak areas were normalized to the appropriate internal standard peak area; d_0_-ω-OH internal standards were used to quantify any oxidized d_7_-C12- or d_7_-C16-BAC metabolite that is not a COOH product; d_0_-COOH BACs were used to quantify d_7_-COOH BAC products; and d_0_-BACs were used to quantify the d_7_-BAC parent levels. All data are presented as the median. Statistical analyses were performed on GraphPad Prism (GraphPad Software, La Jolla, CA) using Welch’s one-way ANOVA analysis, followed by Dunnett’s multiple comparison tests relative to the Control.

## Results

To assess the consequences of oral BAC exposure on the gut microbiome, C57BL/6J male and female mice were exposed to either control Nutra-Gel diet or Nutra-Gel diet with added d_7_-C12-BAC or d_7_-C16-BAC at a dosage of 120 µg/g/day for a duration of 1-week. This dosage paradigm (Figure 2) was adapted from Melin et al., who had previously revealed that 120 µg/g/day was half of the lowest observable adverse effect limit (LOAEL) (Melin *et al*., 2014).

**Figure 2.**
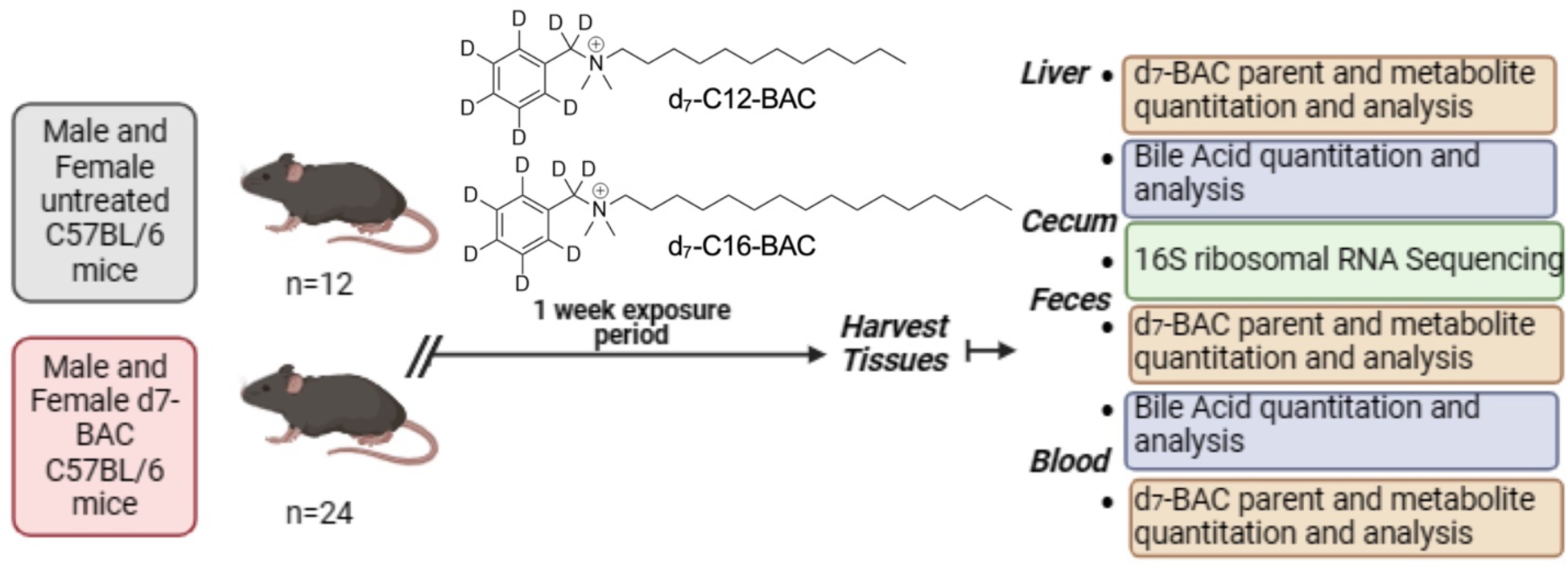
Schematic overview of the experimental design. Male and female C57BL/6 mice were randomly assigned to either the control group, or d_7_-C12- or d_7_-C16-BAC exposed group; each had N=6. Mice were exposed via a gel diet for one week. Harvested tissues were utilized for either d_7_-BAC parent and metabolite analysis, bile acid analysis, or 16S ribosomal RNA sequencing, as described in *Materials and Methods*.

### BAC and BAC metabolite distribution

BACs were quantified in male and female liver, feces, and blood (Figures 3A and 3B). Neither d_7_-C12-BAC nor d_7_-C16-BAC was detected in mice fed the control diet. The highest levels of BACs were observed in the fecal samples, reaching 100s of μM to low mM concentrations, while their levels were in a few to 100s nM range in the liver and blood. Comparing the two sexes, higher levels of parent C12- and C16-BACs were generally observed in female samples, particularly in blood and feces. Metabolites of BACs are formed by cytochrome P450 oxidation of the terminal (ω and ω-1) carbon atoms on the alkyl chain in the parent BAC structure (Seguin *et al*., 2019a). Quantifiable levels of ω- and (ω-1)-hydroxy (OH) metabolites of both d_7_-C12- and d_7_-C16-BAC structures were observed in the respective exposed group (Figures 3A and 3B). Furthermore, products from β-oxidation of the carboxylic acid (COOH) metabolites were observed in male and female blood, liver, and feces. Even numbered chain-shortened carboxylic acid BACs (C6, C8, C10, C12, C14, and C16) were all quantified in the blood, liver, and feces of both d_7_-C12- and d_7_-C16-BAC exposed groups (Figure 4B-E), with the feces containing the highest levels of individual COOH BACs (up to mM), followed by livers (up to 1000s nM) and blood (generally < 100 nM). Again, female samples appear to contain higher levels of COOH BACs than male samples, particularly in the feces. Carboxylic acid metabolites, d_7_-COOH C14-BAC and d_7_-COOH C16-BAC, products of d_7_-C16-BAC, were only seen in d_7_-C16-BAC-treated mice. We also observed a series of odd-chain carboxylic acids but at much lower levels (Supplemental Tables 6 and 7). The initial odd-chain carboxylic acid could be formed from terminal C-C cleavage mediated by cytochromes P450. Such reactions are known in other CYP-mediated metabolisms (Guengerich and Yoshimoto, 2018a; Yoshimoto *et al*., 2016a; Stok and De Voss, 2000a; Cryle and De Voss, 2004a; Umehara, Kudo, Hirao, Morita, Uchida, *et al*., 2000a; Umehara, Kudo, Hirao, Morita, Ohtani, *et al*., 2000a; Umehara *et al*., 2004a), but the specific CYP isoform responsible for the C-C cleavage of BACs remains to be elucidated.

**Figure 3.**
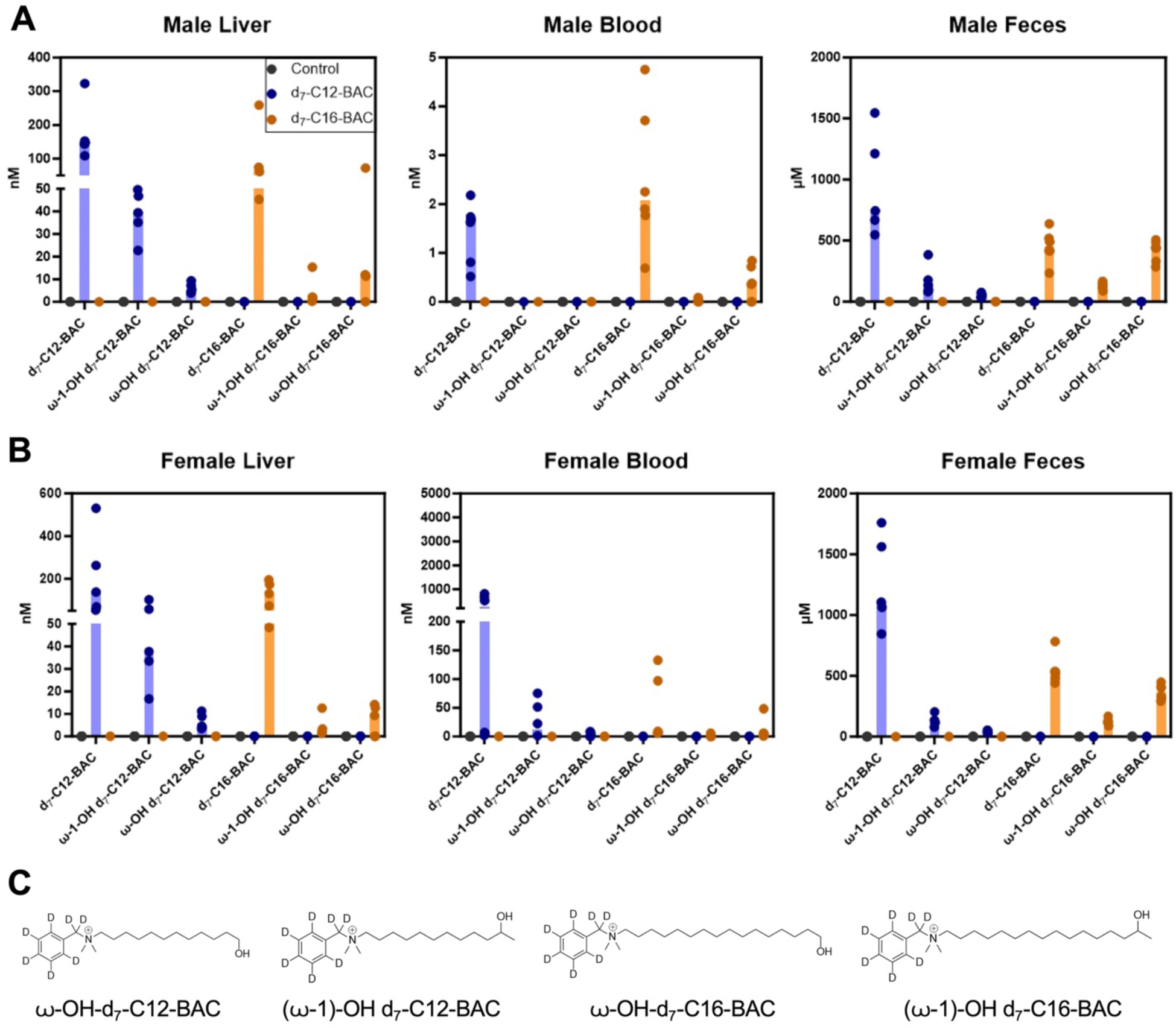
Quantified levels of parent BACs and their hydroxy metabolites in Male **(A)** and Female **(B)** liver, blood, and feces. Male and female C57BL/6 mice were randomly assigned to either the control group, (120 µg/g/day) d_7_-C12-BAC group, or (120 µg/g/day) d_7_-C16-BAC group; each had N=6. Mice were exposed via a gel diet for one week. C. Chemical structures of (ω-OH)-d_7_-C12-BAC, (ω-1)-OH d_7_-C12-BAC, (ω-OH)-d_7_-C16-BAC and (ω-1)-OH-d_7_-C16-BAC.

**Figure 4.**
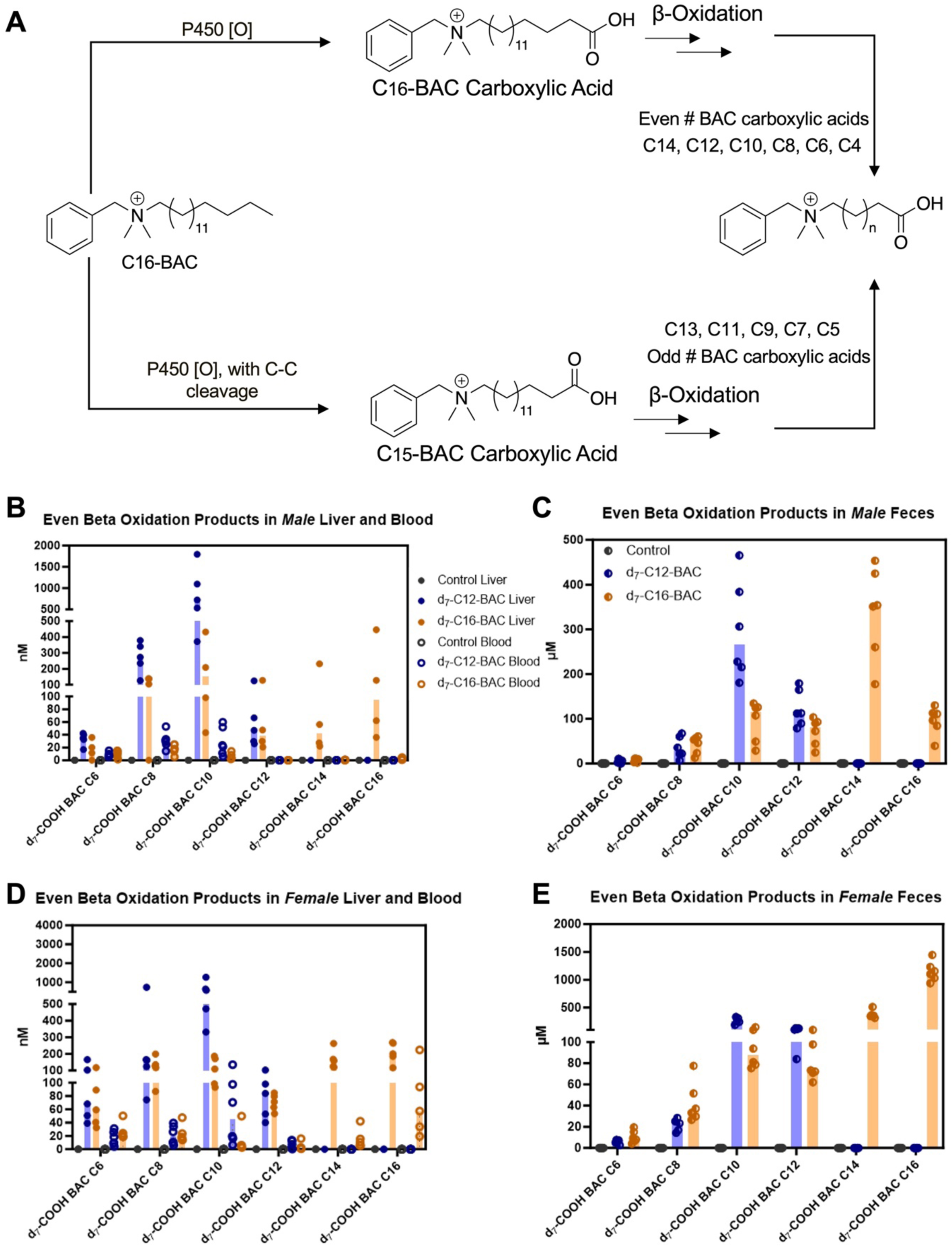
Formation and detection of beta-oxidation products from BACs. A. Schematic overview of proposed pathways of the formation of beta-oxidation products. B-E. Even beta-oxidation products were quantified in Male (B and C) and Female (D and E) liver, blood, and feces using targeted mass spectrometry as described in *Materials and Methods*. N=4-6 per group.

### Effects of BACs on gut microbiome using 16S rRNA gene sequencing

To evaluate the effects of BAC exposure on the gut microbiome, 16S rRNA sequencing targeting the V4 region was performed in the cecum of male and female C57BL/6J mice exposed to either control, d_7_-C12-BAC (120 µg/g/day), or d_7_-C16-BAC (120 µg/g/day). Alpha diversity, a description of species richness, was assessed by the Rarefaction Plot on QIIME2. In both male and female cohorts, observed features for any given sequencing depth were lower in BAC-treated mice than in controls, indicating a decrease in microbial richness compared to control mice (Figure 5A-B). Beta diversity, a measurement of species differences amongst samples, was determined and visualized by principal coordinates analysis (PCoA) using Emperor (Vázquez-Baeza *et al*., 2013a). The female cohort had a clear separation of treatment groups, indicating distinct microbial communities amongst the controls, d_7_-C12-, and d_7_-C16-BAC female mice. In the male cohort, while the separation was apparent between the BAC-treated mice and controls, large variability in the d_7_-C16-BAC group led to overlaps with the d_7_-C12-BAC group (Figure 5C-D).

**Figure 5.**
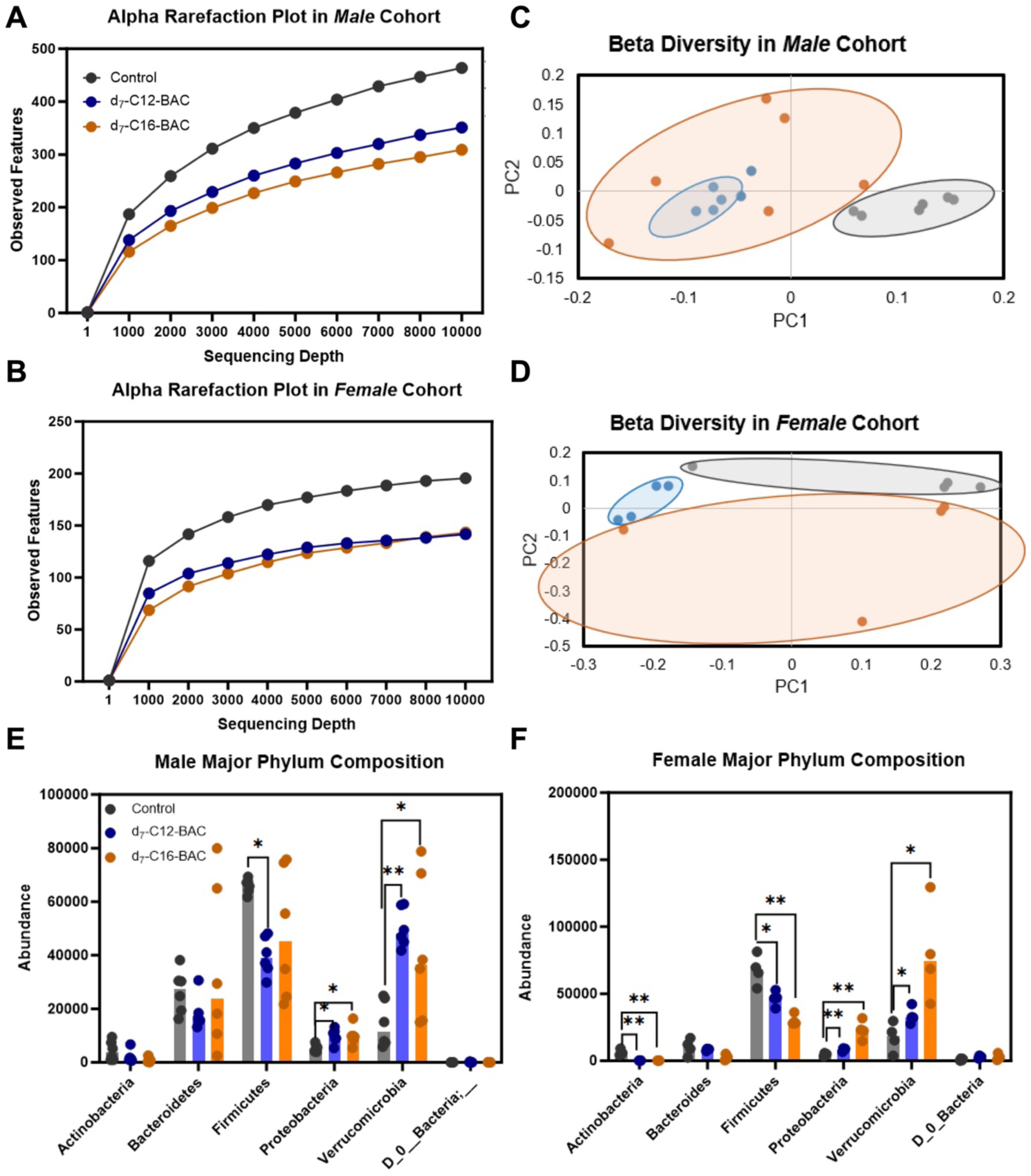
Diversity and Phyla Taxonomic Analysis. **A, B.** Rarefaction plots demonstrating Alpha Diversity in Male (A) and Female (B) cohorts. **C, D.** Beta diversity PCoA plots were plotted via weighted unifrac distance matrix on qiime2. PC1 and PC2 are plotted for both Male and Female cohorts (*Materials and Methods*). **E, F.** Major Phylum compositions of Male (E) and Female (F) cohorts. Data are presented as median; asterisks signify statistical significance by one-way ANOVA using Welch’s test. Comparisons are relative to control. Male (N = 6 per group), Female (N=4 per group). * *P* ≤ 0.05; ** *P* ≤ 0.01; *** *P* ≤ 0.001; **** *P* ≤ 0.0001.

Taxa analysis by QIIME2 revealed the composition of the gut microbiome in different groups: control, d_7_-C12-BAC, and d_7_-C16-BAC exposed mice (Figure 5E-F). The majority of operational taxonomic units were assigned to phyla Firmicutes (41.8% in females; 40.8% in males), Verrucomicrobia (38.0% in females, 28.4% in males), Bacteroides (6.1% in females, 21.1% in males), Proteobacteria (10.4% in females; 6.7% in males), and Actinobacteria (1.8% in females; 1.9% in males). 1.8% in females and 0.05% of all defined taxonomical units in the males were not assigned to any bacteria phylum. Actinobacteria phyla were greatly decreased by both d_7_-C12-BAC and d_7_-C16-BAC treatments in the female cohort. Both Proteobacteria and Verrucomicrobia were significantly increased in d_7_-C12-BAC- and d_7_-C16-BAC-treated mice in both male and female cohorts, relative to controls. Notably, Firmicutes was significantly lower in both BAC-exposed groups in the female cohort but only in the d_7_-C12-BAC group of the male cohort. Bacteroides was not significantly changed following BAC exposure in either male or female cohorts.

Analysis at the family level revealed identifications of taxa belonging to the Firmicutes, Verrucomicrobia, Proteobacteria, Bacteroides, and Actinobacteria phyla (Figure 6A-B). The composition of family microbial taxa was changed in BAC-treated mice compared to controls. The sums of individual counts of family taxa were plotted and not found to be significantly different between treatment groups, indicating that shifts in family microbial taxa were not the result of lowered or higher counts of family microbial taxa. The top 10 abundant families of bacteria are plotted via heat maps. Notably, Akkermansiaceae, of the Verrucomicrobia phylum, was the most abundant family in both male and female cohorts (28.2 and 38.0%). Additionally, BAC exposure led to increased Akkermansiaceae in both male and female cohorts. Lachnospiraceae and Ruminococcaceae were also notable contributors to the family microbial composition, and both families were significantly lower in the composition of the BAC-exposed male and female mice. Burkholderiaceae (of the Proteobacteria phylum) and Muribaculaceae (of the Bacteroides phylum) were both significantly different in abundance in BAC-exposed mice relative to controls.

**Figure 6.**
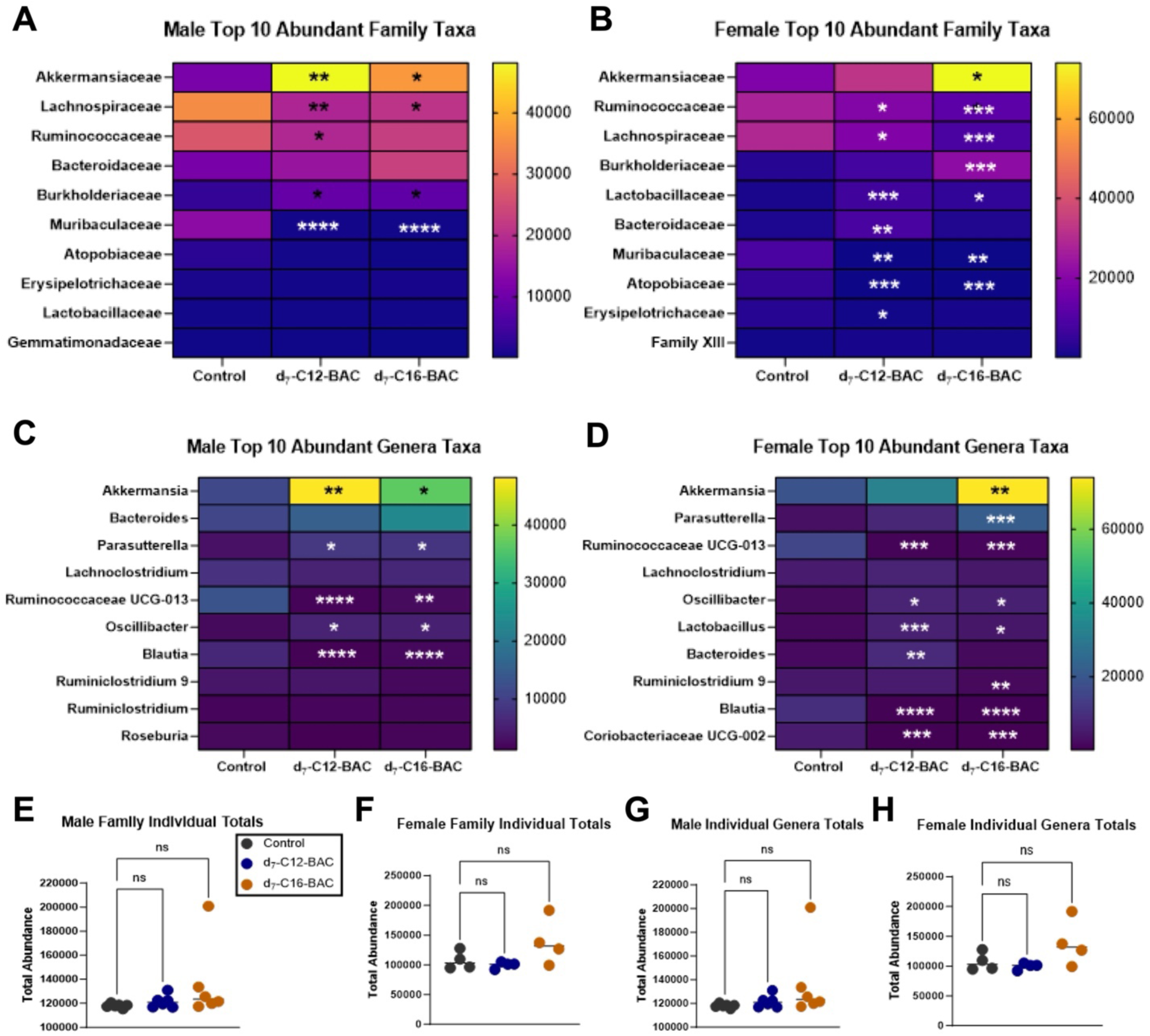
Taxonomic Analysis at the Family and Genera Levels. **A, B.** Heatmaps demonstrating the top 10 abundant families of bacteria in Male (A) and Female (B) cohorts. **C, D.** Heatmaps demonstrating the top 10 abundant genera of bacteria detected in Male (C) and Female (D) cohorts. Each column in heatmaps (A-D) showcases the median of the respective group: control, d_7_-C12-BAC, and d_7_-C16-BAC. Taxonomic analysis was completed as described in materials and methods by Qiime2. **E-H**, Sums of either family or genera totals per sample were plotted and assessed for statistical significance in Male (E, G) and Female (F, H) cohorts. All data are presented as medians; asterisks signify statistical significance by one-way ANOVA using Welch’s test. Comparisons are relative to control. Male (N=6 per group), Female (N=4 per group). * *P* ≤ 0.05; ** *P* ≤ 0.01; *** *P* ≤ 0.001; **** *P* ≤ 0.0001.

Analysis at the genus level revealed the identification of taxa primarily in Firmicutes phyla (Figure 6C-D). In the female cohort, six of the top ten most abundantly identified genera belong to the Firmicutes phyla, including Ruminococcaceae UCG-013, Lachnoclostridium, Oscillibacter, Lactobacillus, Ruminiclostridium 9, and Blautia. Blautia and Ruminococcaceae UCG-013 were significantly decreased in d_7_-C12-BAC and d_7_-C16-BAC exposed mice, while Lactobacillus was significantly increased. Additionally, chain length-specific alterations occurred; Oscillibacter was only significantly increased in d_7_-C12-BAC-exposed female mice, and Ruminiclostridium 9 was only significantly decreased in d_7_-C16-BAC-exposed female mice. In the male cohort, six of the top ten most abundantly identified genera belong to the Firmicutes phyla as well: Lachnoclostridium, Ruminococcaceae UCG-013, Oscillibacter, Ruminiclostridium 9, Blautia and Roseburia. Notably, Oscillibacter, Parasutterella, and Akkermansia were significantly increased in d_7_-BAC-exposed male mice relative to controls. Blautia and Ruminococcaceae UCG-013 were significantly lowered in d_7_-BAC-exposed male mice relative to controls. Akkermansia genera described the most significant proportion of the identified genera in the microbiomes of both the male and female cohorts. Additionally, Coriobacteriaceae UCG-002, part of Actinobacteria phyla, was identified within the top 10 most abundant genera in the female cohort and was significantly lower in BAC-treated female mice relative to controls. Total individual genera count within the treatment groups showed no significant difference in male and female cohorts, indicating that changes noted in the composition are not due to a loss or gain of microbiota within a treatment group.

### BAs in feces and liver

To understand how oral BAC exposure impacts BA homeostasis, a targeted BA quantitation assay was developed based on previous methods (see Materials and Methods) and performed on the blood, liver, and feces of male and female C57BL/6J mice exposed to either control, d_7_-C12-BAC, or d_7_-C16-BAC (120 µg/g/day) (n = 4-6 per group).

Primary BAs, including cholic acid (CA), chenodeoxycholic acid (CDCA), α-muricholic acid (α-MCA), and β-muricholic acid (β-MCA), were quantified in the male and female liver (Figure 7A) and fecal (Figure 7B) extracts. In the female and male liver extracts, primary bile acids, CA, CDCA, α-MCA, and β-MCA amounts were not significantly altered in d_7_-C12- or d_7_-C16-BAC treated mice relative to controls. Notably, in the female and male fecal extracts, the d_7_-C16-BAC treated mice had a significantly higher level of CA, a product of cholesterol, than controls.

**Figure 7.**
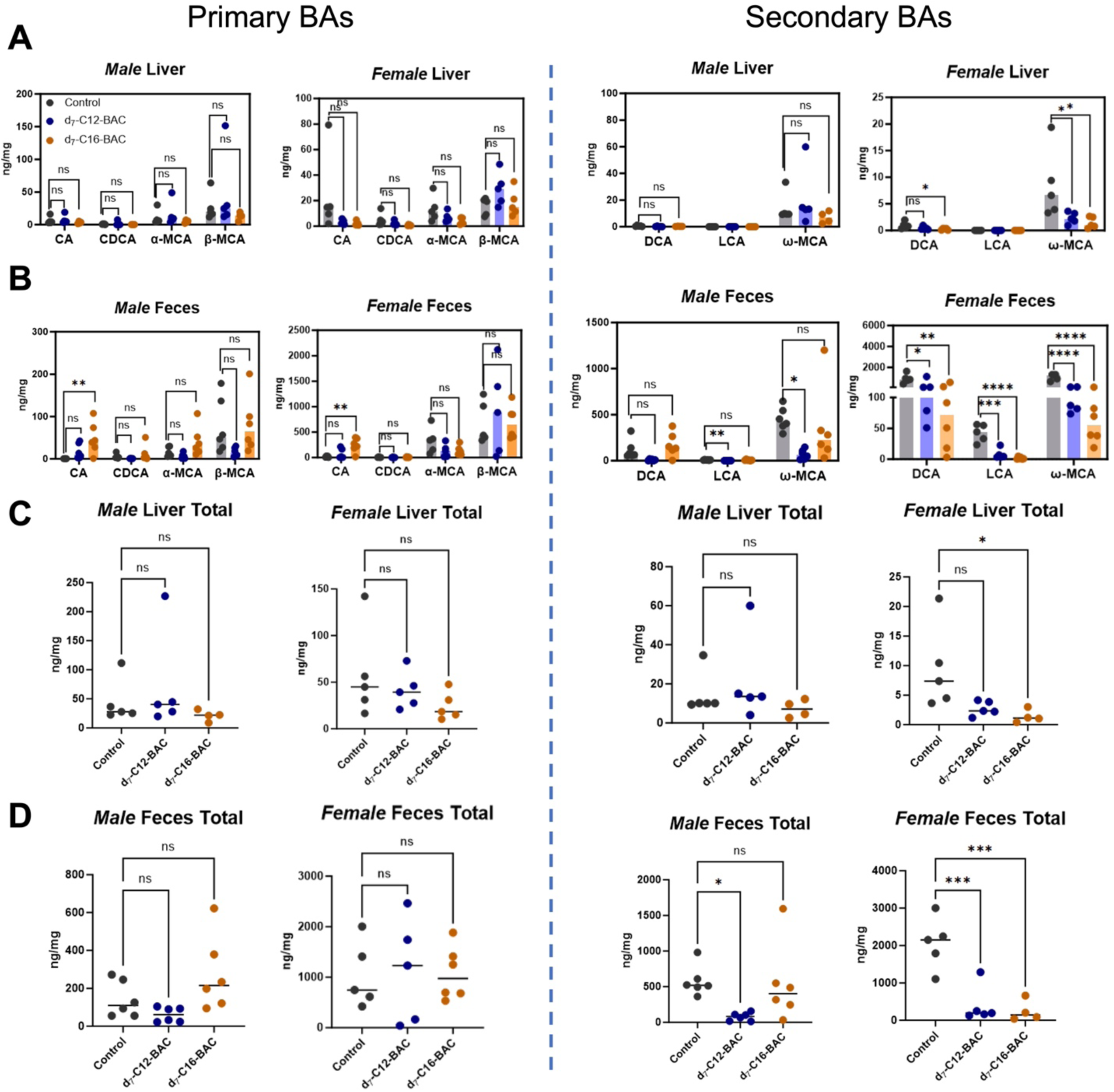
Quantified primary and secondary bile acids in Male and Female mice. **A.** Primary (CA, CDCA, α-MCA, and β-MCA) and secondary (DCA, LCA, ω-MCA) BAs, quantified in liver extracts of control, d_7_-C12- and d_7_-C16-BAC (120 µg/g/day) male and female mice. N=4-6. B. Primary and Secondary BAs quantified in feces of control, d_7_-C12-BAC, and d_7_-C16-BAC (120 µg/g/day) male and female mice. N=5-6. **C-D.** Sums of either primary or secondary bile acids in male and female liver (C) and feces (D). All data are presented as medians; asterisks signify statistical significance by one-way ANOVA using Welch’s test. * *P* ≤ 0.05; ** *P* ≤ 0.01; *** *P* ≤ 0.001; **** *P* ≤ 0.0001.

Secondary bile acids, DCA, LCA, and ω-MCA, were also quantified in liver and fecal extracts (Figure 7A and 7B). Notably, in the female liver, both DCA and ω-MCA were significantly lower in d_7_-C16-BAC treated mice relative to controls. ω-MCA was also significantly lower in the d_7_-C12-BAC female liver. There were no significant differences in DCA or ω-MCA amounts in the three treatment groups in the male liver. LCA was not detected in either male or female liver. Significantly, d_7_-C12-BAC and d_7_-C16-BAC led to greatly decreased amounts of secondary BAs, DCA, LCA, and ω-MCA in female feces. In the male cohort, only d_7_-C12-BAC treated mice had statistically lower amounts of LCA and ω-MCA in feces.

Regulation of the BA pool was further analyzed by summing the total BAs (either primary or secondary) per sample in the liver and feces (Figure 7 Panel C, Panel D). Total primary BA content was not significantly different between d_7_-BAC exposed mice and controls in feces or liver in male and female cohorts. However, total secondary BA content was significantly lower in the d_7_-C16-BAC exposed female livers, but not male livers, relative to controls. In the feces of both male and female d_7_-C12-BAC exposed mice, total secondary BA content was significantly lower. d_7_-C16-BAC treated mice also had significantly lowered total secondary BA content in the feces of the female cohort, but not the male cohort, relative to controls.

## Discussion

The gut microbiome has become increasingly recognized as an essential marker for numerous disease pathologies (Chiang and Lin, 2019; Manos, 2022; Wu and Lewis, 2013; Wang *et al*., 2021; Boursier and Diehl, 2015). Various groups have studied the potential of environmental toxicant exposure on microbiome composition. For example, cadmium exposure altered energy metabolism and gut microbiota composition in male C57BL/6J mice (Zhang *et al*., 2015). Li et al. found that environmental contaminants, polybrominated diphenyl ethers, altered gut microbiome composition and bile acid homeostasis in male C57BL/6J mice (Li *et al*., 2018). Understanding the mechanisms by which environmental agents can induce changes in the gut-liver axis is important in elucidating the association between environmental exposure and human health.

Alpha diversity has been studied extensively and is significantly decreased in subjects in various metabolic disease states (Carroll *et al*., 2012; Shen *et al*., 2017; Palmas *et al*., 2021). Alpha diversity analysis demonstrates that exposure to either d_7_-C12-BAC or d_7_-C16-BAC led to decreased microbial richness in both male and female cohorts (Figure 2A, 2B). Beta diversity analysis shows that the microbial communities differ between the BAC-treated and control mice in male and female cohorts (Figure 2C, 2D).

Our results are the first to show BAC exposure can alter the gut microbiota composition. We observed that gram-negative phyla (Proteobacteria and Verrucomicrobia) were significantly increased in BAC-treated mice relative to controls (Figure 5E-F). In contrast, the gram-positive phylum (Firmicutes and Actinobacteria) was significantly decreased relative to controls. In human populations, the predominant phyla are Firmicutes, Bacteroidetes, Actinobacteria, and Proteobacteria (Rizzatti *et al*., 2017).

The Firmicutes phylum has essential roles in the fermentation of dietary fibers, producing metabolites like vitamins and short-chain fatty acids. Because BAC-treated mice have lowered Firmicutes relative to controls, a consequence could be less short-chain fatty acid formation. Less short-chain fatty acid formation can result in effects like inflammation or deprived gut lining cells. In male and female cohorts, the Proteobacteria phylum was significantly increased in both d_7_-C12-BAC and d_7_-C16-BAC treated mice relative to controls. Proteobacteria is often increased in disease and has been noted as a potential marker of microbiota instability and predisposition of inflammatory-sustained disease onset (Rizzatti *et al*., 2017). The consistent increases in Proteobacteria in BAC-exposed mice could foreshadow subsequent proinflammatory diseases.

Notably, the Actinobacteria phylum was significantly decreased in BAC-treated female mice relative to controls (Figure 5F). Actinobacteria are recognized contributors to maintaining gut barrier homeostasis and are also involved in the transformation of linoleic acid into conjugated linoleic acids, which have health-promoting effects like anti-obesity and anti-diabetes (Binda *et al*., 2018). The Coriobacteriia class was the predominant bacteria belonging to the Actinobacteria phylum in both cohorts. While the effects of BAC exposure on Actinobacteria phylum are notable in the female cohort, the male cohort did not show significant changes. This difference in the microbiome changes between the male and female cohort could result from the variability in the male d_7_-C16-BAC group, and such variability could also explain the nuances in secondary bile acid formation.

Our study also demonstrates BAC exposure led to compositional shifts in family taxa (Figure 6A-B). Absolute total microbial counts within each treatment group were not significantly different (Figure 6E-F), indicating that the differences in the abundance of bacteria are results of compositional shifts. Notably, in both d_7_-C12-BAC and d_7_-C16-BAC male mice and d_7_-C16-BAC female mice, Akkermansiacae and Akkermansia of the Verrucomicrobia phyla were significantly increased. Akkermansia is currently the only defined genus within the Akkermansiaceae family and is known to specialize in mucin degradation. Akkermansia is consistently recognized for its beneficial role in maintaining a healthy mucosal layer and gut barrier integrity (González *et al*., 2023). Interestingly, Akkermansia decreases in various disease states and metabolic disorders like inflammatory bowel disease, obesity, and diabetes (González *et al*., 2023).

Lachnospiraceae and Ruminococcaceae of the Firmicutes phylum decreased in d_7_-C12- and d_7_-C16-BAC treated mice relative to controls. Lachnospiraceae and Ruminococcaceae have roles in secondary bile acid formation. Thus, their decreases could be correlated to the lowered secondary bile acid quantitation in male and female feces. Atopobiaceae, a part of the Actinobacteria phylum, was significantly decreased in female d_7_-C12- and d_7_-C16-BAC treated mice. Atopobiaceae is lactate-producing and, when isolated from mouse intestines, showed high resistance to mammalian bile extracts due to their significant bile salt hydrolase activity. This finding is significant in the context of differences between the male and female cohort observations regarding secondary bile acid formation (Morinaga *et al*., 2022).

Our study also identifies specific genera that were impacted by oral BAC exposure in male and female cohorts (Figure 6C-D). Parasutterella was significantly higher in both male d_7_-C12-BAC and d_7_-C16-BAC-treated mice and female d_7_-C16-BAC-treated mice. Parasutterella is supportive of bile acid maintenance and cholesterol metabolism. Parasutterella isolates are asaccharolytic and succinate producers (Ju *et al*., 2019). BAC exposure leading to Parasutterella increases could be evidence of the microbiota trying to mediate the decreases of other important bacteria in bile acid maintenance, like Lachnospiraceae, Lactobacilliceae, and Ruminococcaceae.

Lastly, in both male and female cohorts, Blautia was significantly decreased in both d_7_-C12- and d_7_-C16-BAC-treated mice relative to controls. Blautia has become extensively recognized as a genus capable of using glucose for carbohydrate utilization. Different strains demonstrate varying capacities to use sucrose, fructose, lactose, and more. Blautia can ferment glucose and produce final products, acetic acid, succinic acid, lactic acid, and ethanol. Notably, certain Blatuia species can perform 7ɑ-dehydroxylation of primary bile acids and convert them to secondary bile acids like lithocholic and deoxycholic acid (Liu *et al*., 2021). Lactobacillus was increased in both d_7_-C12- and d_7_-C16-BAC female groups. Various Lactobacillus species have been recognized for their ability to deconjugate bile acids in the GI tract through BSH proteins (O’Flaherty *et al*., 2018). Ruminiclostridium 9, of the Firmicutes phyla was significantly decreased in the d_7_-C16-BAC group relative to female controls and has been reported to be a short-chain fatty acid (SCFA) producer in the gut (Zhao *et al*., 2022). Lastly, in the female cohort only, both d_7_-C12- and d_7_-C16-BAC led to Coriobacteriaceae UCG-002 of the Actinobacteria phyla being significantly decreased. Coriobacteriaceae UCG-002 has been reported to be positively correlated with serum triglycerides, total cholesterol, and LDL cholesterol levels, as well as serum TNF-a, IL-6, and LPS levels, suggesting a significant role in the female immune response. Other work has shown the correlation between Coriobacteriaceae UCG-002 and SCFAs (Pradista *et al*., 2022).

Our study also describes the metabolic route of BAC structures in an orally exposed mouse model. Our previous work elucidated the cytochrome P450-mediated oxidation of BACs, which occurs on the alkyl chain region (Seguin *et al*., 2019), forming primary ⍵- and (⍵-1)-hydroxy-BAC metabolites and secondary ⍵-COOH-BAC metabolite. Both ⍵- and ⍵-1 oxidized metabolites of deuterated d_7_-C12- and d_7_-C16-BAC were quantified in male and female liver, blood, and feces. Furthermore, we observed evidence that BAC COOH metabolites formed from CYP-mediated metabolism underwent further metabolism by β-oxidation. Most CYP oxidation of BACs occurs on the even-numbered alkyl chain, such as those of the d_7_-C12- or d_7_-C16-BAC structures, to produce even-numbered alkyl chain COOH metabolites. However, if CYP oxidation catalyzes a carbon-carbon cleavage, the resulting removal of the ω-carbon leaves β-oxidation to occur on an odd-numbered alkyl chain (Umehara, Kudo, Hirao, Morita, Uchida, *et al*., 2000a; Umehara, Kudo, Hirao, Morita, Ohtani, *et al*., 2000a; Umehara *et al*., 2004a) , and subsequent chain-shortened COOH BAC products will contain an odd-numbered carbon alkyl chain length. The levels of odd-numbered β-oxidation products of d_7_-C12- and d_7_-C16-BAC in male and female blood, liver, and feces were much lower than that of the even-numbered β-oxidation products, indicating that P450 oxidation occurring with a carbon-carbon cleavage is a minor pathway (Supplemental Tables S6-S7 and Figures S1-S4).

In conclusion, our study provides novel insights into benzalkonium chloride toxicity and metabolism following an oral dose model. We showed that BACs decreased microbial richness and elicited a microbial community different than control mice in a sex-dependent manner. The changes in the microbial community include bacteria related to bile acids, gut barrier homeostasis, short-chain fatty acids, and pro-inflammatory biomarkers. Notably, we observed decreased levels of secondary bile acid-forming bacteria, which may be responsible for the lowered levels of secondary BAs in the feces of both sexes and in the female liver. Remarkably, we observed different microbial communities between male and female cohorts. We believe these differences discussed may be responsible for secondary BAs decreasing much more in female fecal extracts than the males. Additionally, quantitative analysis of BAC metabolites in the liver, blood, and feces provided evidence of β-oxidation as a major metabolic route following CYP-mediated oxidation. This present study is the first to elucidate the capacity of ubiquitously used BACs to alter an emerging health biomarker, the gut microbiome, and the gut-liver axis. Future work should consider other gut microbiome axes and clinically relevant lifetime exposure to either occupational exposure or demographic exposure to BACs.

## Supporting information

Supplemental 16S sequencing data

