## Supplemental Tables and Figures for "Oral Exposure to Benzalkonium Chlorides in Male and Female Mice Reveals Sex-Dependent Alteration of the Gut Microbiome and Bile Acid Profile"

Libin Xu, PhD

University of Washington, Box 357610

H172 Health Science Building

Seattle, WA 98195-7610

| Table of Contents | |
|---|---|
| Supplemental Table S1. Sciex 6500 conditions for bile acid (BA) Analysis | Page 4 |
| Supplemental Table S2. MRM Transition Conditions for BA Analysis | Page 5 |
| Supplemental Table S3. 16S rRNA raw Paired End (PE) sequencing reads | Pages 6-7 |
| Supplemental Table S4. Average and Standard Deviations (Stdev) of parent BACs, hydroxylated metabolites, and even chained beta-oxidation products in Male and Female Liver and Blood. | Pages 8-10 |
| Supplemental Table S5. Average and Standard Deviations (Stdev) of parent BACs, hydroxylated metabolites, and even chained beta-oxidation products in Male and Female Feces. | Pages 11-12 |
| Supplemental Table S6. Average and Standard Deviations (Stdev) of odd chained beta-oxidation products in Male and Female Liver and Blood. | Pages 13-14 |
| Figure S1. Odd Chained Beta Oxidation Products in Male Liver and Blood | Page 15 |

| | |
|---|---|
| Figure S2. Odd Chained Beta Oxidation Products in Female Liver and Blood | Page 16 |
| Supplemental Table S7. Average and Standard Deviations (Stdev) of odd chained beta-oxidation products in Male and Female Feces. | Page 17 |
| Figure S3. Odd Chained Beta Oxidation Products in Male Feces | Page 18 |
| Figure S4. Odd Chained Beta Oxidation Products in Female Feces | Page 19 |
| Supplemental Table S8. Precursor and Product Ion Transitions for BAC Quantitation | Pages 20-22 |

| Supplemental Table S1. Sciex 6500 conditions for BA Analysis | |
|---|---|
| *Source/Gas* | |
| Ion Source | Turbo Spray Ion Drive |
| Curtain Gas (CUR) | 40 |
| Collision Gas (CAD) | 12 |
| Ion Spray Voltage (IS) | -4500 |
| Temperature (TEM) | 450 |
| Ion Source Gas 1 (GS1) | 50 |
| Ion Source Gas 2 (GS2) | 70 |
| *UPLC Sample Manager* | |
| Column Temperature | 45°C |
| Sample Temperature | 5°C |

**Supplemental Table S2.** Multiple Reaction Monitoring (MRM) transition conditions in both positive and negative mode. Abbreviations: DP = Declustering Potential; EP = Entrance Potential; CE = Collision Energy; CXP = Collision Cell Exit Potential; Q = Quadrupole Mass Filter

| **Supplemental Table 2. MRM Transition Conditions for BA Analysis** | | | | | | |
|---|---|---|---|---|---|---|
| Negative Mode | | | | | | |
| Bile Acid | Q1 | Q3 | DP (volts) | EP (volts) | CE (volts) | CXP (volts) |
| d0-α-MCA | 407.2 | 387.2 | -200 | -10 | -48 | -19 |
| d0-β-MCA | 407.2 | 371.3 | -210 | -10 | -44 | -19 |
| d0-ω-MCA | 407.2 | 387.1 | -195 | -10 | -46 | -23 |
| d0-CA | 407.2 | 343.1 | -210 | -10 | -44 | -23 |
| d0-CDCA | 391.2 | 373.2 | -185 | -10 | -44 | -19 |
| d0-DCA | 391.2 | 345.2 | -180 | -10 | -46 | -23 |
| d4-α-MCA | 411.2 | 390.1 | -200 | -10 | -48 | -21 |
| d4-ω-MCA | 411.2 | 390 | -200 | -10 | -48 | -23 |
| d4-CA | 411.2 | 347.2 | -150 | -10 | -46 | -21 |
| d4-DCA | 395.2 | 349.2 | -175 | -10 | -46 | -21 |
| Positive Mode | | | | | | |
| d0-CA | 359.2 | 135.1 | 130 | 10 | 35 | 10 |
| d4-LCA | 363.2 | 135.1 | 130 | 10 | 35 | 10 |

**Supplemental Table S3**. 16S rRNA raw Paired End (PE) sequencing reads for cecal samples from C57BL/6 male and female mice. Control, $d_7$-C12-BAC (120 µg/g/day), $d_7$-C16-BAC (120 µg/g/day). n=4-6 per group.

| Sample Name | Raw PE Reads |
|---|---|
| Male Control_1 | 134160 |
| Male Control_2 | 134526 |
| Male Control_3 | 135380 |
| Male Control_4 | 134794 |
| Male Control_5 | 132596 |
| Male Control_6 | 136532 |
| Male $d_7$-C12-BAC_1 | 133989 |
| Male $d_7$-C12-BAC_2 | 135309 |
| Male $d_7$-C12-BAC_3 | 134357 |
| Male $d_7$-C12-BAC_4 | 132793 |
| Male $d_7$-C12-BAC_5 | 145379 |
| Male $d_7$-C12-BAC_6 | 138315 |
| Male $d_7$-C16-BAC_1 | 135747 |
| Male $d_7$-C16-BAC_2 | 142259 |
| Male $d_7$-C16-BAC_3 | 228443 |
| Male $d_7$-C16-BAC_4 | 136267 |
| Male $d_7$-C16-BAC_5 | 144825 |
| Male $d_7$-C16-BAC_6 | 135997 |
| Female Control_1 | 144,957 |
| Female Control_2 | 125,727 |
| Female Control_3 | 107,675 |
| Female Control_4 | 121,833 |

| | |
|---|---|
| Female $d_7$-C12-BAC_1 | 113,114 |
| Female $d_7$-C12-BAC_2 | 103,262 |
| Female $d_7$-C12-BAC_3 | 116,317 |
| Female $d_7$-C12-BAC_4 | 113,081 |
| Female $d_7$-C16-BAC_1 | 109,963 |
| Female $d_7$-C16-BAC_2 | 212,001 |
| Female $d_7$-C16-BAC_3 | 140,080 |
| Female $d_7$-C16-BAC_4 | 151,456 |

**Supplemental Table S4.** Average and Standard Deviations (Stdev) of parent BACs, hydroxylated metabolites, and even chained beta-oxidation products in Male and Female Liver and Blood.

| Male Liver (nM) | | | | | | |
|---|---|---|---|---|---|---|
| *Analyte* | *Control Average* | *Control Stdev* | *$d_7$-C12-BAC Average* | *$d_7$-C12-BAC Stdev* | *$d_7$-C16-BAC Average* | *$d_7$-C16-BAC Stdev* |
| $d_7$-C12-BAC | 0.00 | 0.00 | 174.64 | 84.44 | 0.00 | 0.00 |
| $d_7$-C16-BAC | 0.00 | 0.00 | 0.00 | 0.00 | 109.92 | 99.76 |
| ω-1 OH-$d_7$-C12-BAC | 0.00 | 0.00 | 38.76 | 10.63 | 0.00 | 0.00 |
| ω-OH-$d_7$-C12-BAC | 0.00 | 0.00 | 6.08 | 2.22 | 0.00 | 0.00 |
| ω-1 OH-$d_7$-C16-BAC | 0.00 | 0.00 | 0.00 | 0.00 | 4.87 | 7.05 |
| ω-OH-$d_7$-C16-BAC | 0.00 | 0.00 | 0.00 | 0.00 | 23.91 | 32.72 |
| $d_7$-COOH BAC C6 | 0.00 | 0.00 | 33.19 | 9.94 | 16.87 | 15.08 |
| $d_7$-COOH BAC C8 | 0.00 | 0.00 | 270.73 | 98.36 | 95.48 | 65.65 |
| $d_7$-COOH BAC C10 | 0.00 | 0.00 | 903.17 | 567.47 | 195.63 | 172.00 |
| $d_7$-COOH BAC C12 | 0.00 | 0.00 | 58.90 | 40.43 | 56.47 | 48.92 |
| $d_7$-COOH BAC C14 | 0.00 | 0.00 | 0.00 | 0.00 | 84.39 | 99.40 |
| $d_7$-COOH BAC C16 | 0.00 | 0.00 | 0.00 | 0.00 | 167.87 | 188.97 |
| Male Blood (nM) | | | | | | |
| *Analyte* | *Control Average* | *Control Stdev* | *$d_7$-C12-BAC Average* | *$d_7$-C12-BAC Stdev* | *$d_7$-C16-BAC Average* | *$d_7$-C16-BAC Stdev* |
| $d_7$-C12-BAC | 0.00 | 0.00 | 1.43 | 0.63 | 0.00 | 0.00 |
| $d_7$-C16-BAC | 0.00 | 0.00 | 0.00 | 0.00 | 1.89 | 1.63 |
| ω-1-OH-$d_7$-C12-BAC | 0.00 | 0.00 | 0.00 | 0.00 | 0.00 | 0.00 |
| ω-OH-$d_7$-C12- | 0.00 | 0.00 | 0.00 | 0.00 | 0.00 | 0.00 |

| | | | | | | |
|---|---|---|---|---|---|---|
| BAC | | | | | | |
| ω-1-OH-d$_7$-C16-BAC | 0.00 | 0.00 | 0.00 | 0.00 | 0.03 | 0.03 |
| ω-OH-d$_7$-C16-BAC | 0.00 | 0.00 | 0.00 | 0.00 | 0.38 | 0.35 |
| d$_7$-COOH BAC C6 | 0.00 | 0.00 | 8.91 | 3.49 | 9.45 | 5.06 |
| d$_7$-COOH BAC C8 | 0.00 | 0.00 | 31.04 | 12.67 | 17.25 | 7.26 |
| d$_7$-COOH BAC C10 | 0.00 | 0.00 | 29.05 | 21.91 | 5.90 | 4.59 |
| d$_7$-COOH BAC C12 | 0.00 | 0.00 | 0.64 | 0.74 | 0.31 | 0.14 |
| d$_7$-COOH BAC C14 | 0.00 | 0.00 | 0.00 | 0.00 | 0.50 | 0.31 |
| d$_7$-COOH BAC C16 | 0.00 | 0.00 | 0.00 | 0.00 | 2.38 | 1.44 |
| Female Liver (nM) | | | | | | |
| *Analyte* | *Control Average* | *Control Stdev* | *d$_7$-C12-BAC Average* | *d$_7$-C12-BAC Stdev* | *d$_7$-C16-BAC Average* | *d$_7$-C16-BAC Stdev* |
| d$_7$-C12-BAC | 0.00 | 0.00 | 210.72 | 197.07 | 0.00 | 0.00 |
| d$_7$-C16-BAC | 0.00 | 0.00 | 0.00 | 0.00 | 124.13 | 62.78 |
| ω-1-OH-d$_7$-C12-BAC | 0.00 | 0.00 | 49.49 | 32.74 | 0.00 | 0.00 |
| ω-OH-d$_7$-C12-BAC | 0.00 | 0.00 | 6.38 | 3.46 | 0.00 | 0.00 |
| ω-1-OH-d$_7$-C16-BAC | 0.00 | 0.00 | 0.00 | 0.00 | 4.45 | 4.62 |
| ω-OH-d$_7$-C16-BAC | 0.00 | 0.00 | 0.00 | 0.00 | 7.22 | 6.82 |
| d$_7$-COOH BAC C6 | 0.00 | 0.00 | 85.40 | 51.11 | 67.83 | 35.31 |
| d$_7$-COOH BAC C8 | 0.00 | 0.00 | 253.75 | 275.72 | 133.40 | 41.10 |
| d$_7$-COOH BAC C10 | 0.00 | 0.00 | 660.03 | 360.11 | 131.87 | 43.81 |

| | | | | | | |
|---|---|---|---|---|---|---|
| $d_7$-COOH BAC C12 | 0.00 | 0.00 | 75.65 | 27.88 | 70.49 | 12.48 |
| $d_7$-COOH BAC C14 | 0.00 | 0.00 | 0.00 | 0.00 | 167.08 | 57.48 |
| $d_7$-COOH BAC C16 | 0.00 | 0.00 | 0.00 | 0.00 | 207.15 | 62.74 |
| Female Blood (nM) | | | | | | |
| *Analyte* | *Control Average* | *Control Stdev* | *$d_7$-C12-BAC Average* | *$d_7$-C12-BAC Stdev* | *$d_7$-C16-BAC Average* | *$d_7$-C16-BAC Stdev* |
| $d_7$-C12-BAC | 0.00 | 0.00 | 336.61 | 375.01 | 0.00 | 0.00 |
| $d_7$-C16-BAC | 0.00 | 0.00 | 0.00 | 0.00 | 50.29 | 60.34 |
| ω-1-OH-$d_7$-C12-BAC | 0.00 | 0.00 | 25.02 | 31.69 | 0.00 | 0.00 |
| ω-OH-$d_7$-C12-BAC | 0.00 | 0.00 | 2.98 | 3.77 | 0.00 | 0.00 |
| ω-1-OH-$d_7$-C16-BAC | 0.00 | 0.00 | 0.00 | 0.00 | 1.45 | 2.36 |
| ω-OH-$d_7$-C16-BAC | 0.00 | 0.00 | 0.00 | 0.00 | 12.05 | 20.34 |
| $d_7$-COOH BAC C6 | 0.00 | 0.00 | 16.38 | 10.51 | 26.70 | 13.38 |
| $d_7$-COOH BAC C8 | 0.00 | 0.00 | 21.18 | 13.99 | 23.86 | 13.82 |
| $d_7$-COOH BAC C10 | 0.00 | 0.00 | 57.77 | 52.10 | 13.76 | 20.28 |
| $d_7$-COOH BAC C12 | 0.00 | 0.00 | 6.24 | 5.87 | 4.62 | 6.43 |
| $d_7$-COOH BAC C14 | 0.00 | 0.00 | 0.00 | 0.00 | 16.06 | 15.17 |
| $d_7$-COOH BAC C16 | 0.00 | 0.00 | 0.00 | 0.00 | 85.90 | 82.50 |

**Supplemental Table S5.** Average and Standard Deviations (Stdev) of parent BACs, hydroxylated metabolites, and even chained beta-oxidation products in Male and Female Feces.

| Male Feces (μM) | | | | | | |
|---|---|---|---|---|---|---|
| *Analyte* | *Control Average* | *Control Stdev* | *$d_7$-C12-BAC Average* | *$d_7$-C12-BAC Stdev* | *$d_7$-C16-BAC Average* | *$d_7$-C16-BAC Stdev* |
| $d_7$-C12-BAC | 0.00 | 0.00 | 909.55 | 384.93 | 0.00 | 0.00 |
| $d_7$-C16-BAC | 0.00 | 0.00 | 0.00 | 0.00 | 452.43 | 134.39 |
| ω-1-OH-$d_7$-C12-BAC | 0.00 | 0.00 | 174.65 | 110.04 | 0.00 | 0.00 |
| ω-OH-$d_7$-C12-BAC | 0.00 | 0.00 | 50.72 | 17.24 | 0.00 | 0.00 |
| ω-1-OH-$d_7$-C16-BAC | 0.00 | 0.00 | 0.00 | 0.00 | 132.38 | 27.42 |
| ω-OH-$d_7$-C16-BAC | 0.00 | 0.00 | 0.00 | 0.00 | 414.39 | 87.37 |
| $d_7$-COOH BAC C6 | 0.00 | 0.00 | 4.97 | 4.06 | 7.20 | 3.95 |
| $d_7$-COOH BAC C8 | 0.00 | 0.00 | 35.64 | 23.76 | 42.40 | 19.66 |
| $d_7$-COOH BAC C10 | 0.00 | 0.00 | 296.62 | 110.57 | 95.02 | 44.63 |
| $d_7$-COOH BAC C12 | 0.00 | 0.00 | 122.82 | 40.63 | 71.17 | 30.84 |
| $d_7$-COOH BAC C14 | 0.00 | 0.00 | 0.00 | 0.00 | 337.02 | 103.07 |
| $d_7$-COOH BAC C16 | 0.00 | 0.00 | 0.00 | 0.00 | 95.72 | 31.53 |
| Female Feces (μM) | | | | | | |
| *Analyte* | *Control Average* | *Control Stdev* | *$d_7$-C12-BAC Average* | *$d_7$-C12-BAC Stdev* | *$d_7$-C16-BAC Average* | *$d_7$-C16-BAC Stdev* |
| $d_7$-C12-BAC | 0.00 | 0.00 | 1267.53 | 378.79 | 0.00 | 0.00 |
| $d_7$-C16-BAC | 0.00 | 0.00 | 0.00 | 0.00 | 32.92 | 18.14 |
| ω-1-OH-$d_7$-C12-BAC | 0.00 | 0.00 | 128.83 | 46.88 | 0.00 | 0.00 |
| ω-OH-$d_7$-C12- | 0.00 | 0.00 | 45.22 | 9.34 | 0.00 | 0.00 |

| | | | | | | |
|---|---|---|---|---|---|---|
| BAC | | | | | | |
| ω-1-OH-$d_7$-C16-BAC | 0.00 | 0.00 | 0.00 | 0.00 | 122.79 | 26.17 |
| ω-OH-$d_7$-C16-BAC | 0.00 | 0.00 | 0.00 | 0.00 | 367.49 | 61.89 |
| $d_7$-COOH BAC C6 | 0.00 | 0.00 | 5.97 | 2.07 | 10.51 | 5.82 |
| $d_7$-COOH BAC C8 | 0.00 | 0.00 | 21.61 | 5.88 | 42.71 | 19.21 |
| $d_7$-COOH BAC C10 | 0.00 | 0.00 | 266.26 | 56.44 | 96.80 | 27.97 |
| $d_7$-COOH BAC C12 | 0.00 | 0.00 | 113.48 | 19.33 | 79.82 | 16.26 |
| $d_7$-COOH BAC C14 | 0.00 | 0.00 | 0.00 | 0.00 | 374.99 | 70.17 |
| $d_7$-COOH BAC C16 | 0.00 | 0.00 | 0.00 | 0.00 | 1151.02 | 175.65 |

**Supplemental Table S6.** Average and Standard Deviations (Stdev) of odd chained beta-oxidation products in Male and Female Liver and Blood.

| Male Liver (nM) | | | | | | |
|---|---|---|---|---|---|---|
| *Analyte* | *Control Average* | *Control Stdev* | *$d_7$-C12-BAC Average* | *$d_7$-C12-BAC Stdev* | *$d_7$-C16-BAC Average* | *$d_7$-C16-BAC Stdev* |
| $d_7$-COOH BAC C7 | 0.00 | 0.00 | 0.00 | 0.00 | 0.00 | 0.00 |
| $d_7$-COOH BAC C9 | 0.00 | 0.00 | 89.50 | 39.71 | 7.13 | 5.22 |
| $d_7$-COOH BAC C11 | 0.00 | 0.00 | 76.37 | 45.49 | 3.51 | 4.02 |
| $d_7$-COOH BAC C13 | 0.00 | 0.00 | 0.00 | 0.00 | 5.09 | 7.36 |
| $d_7$-COOH BAC C15 | 0.00 | 0.00 | 0.00 | 0.00 | 5.26 | 7.04 |
| Male Blood (nM) | | | | | | |
| *Analyte* | *Control Average* | *Control Stdev* | *$d_7$-C12-BAC Average* | *$d_7$-C12-BAC Stdev* | *$d_7$-C16-BAC Average* | *$d_7$-C16-BAC Stdev* |
| $d_7$-COOH BAC C7 | 0.00 | 0.00 | 2.90 | 0.87 | 0.00 | 0.00 |
| $d_7$-COOH BAC C9 | 0.00 | 0.00 | 5.23 | 2.68 | 0.00 | 0.00 |
| $d_7$-COOH BAC C11 | 0.00 | 0.00 | 9.97 | 12.20 | 0.00 | 0.00 |
| $d_7$-COOH BAC C13 | 0.00 | 0.00 | 0.00 | 0.00 | 0.00 | 0.00 |
| $d_7$-COOH BAC C15 | 0.00 | 0.00 | 0.00 | 0.00 | 0.00 | 0.00 |
| Female Liver (nM) | | | | | | |
| *Analyte* | *Control Average* | *Control Stdev* | *$d_7$-C12-BAC Average* | *$d_7$-C12-BAC Stdev* | *$d_7$-C16-BAC Average* | *$d_7$-C16-BAC Stdev* |
| $d_7$-COOH BAC C7 | 0.00 | 0.00 | 12.35 | 7.64 | 2.81 | 0.81 |
| $d_7$-COOH BAC C9 | 0.00 | 0.00 | 62.46 | 54.59 | 8.16 | 1.28 |
| $d_7$-COOH BAC C11 | 0.00 | 0.00 | 59.72 | 16.65 | 2.34 | 0.65 |

| | | | | | | |
|---|---|---|---|---|---|---|
| $d_7$-COOH BAC C13 | 0.00 | 0.00 | 0.00 | 0.00 | 8.18 | 5.71 |
| $d_7$-COOH BAC C15 | 0.00 | 0.00 | 0.00 | 0.00 | 8.22 | 1.71 |
| Female Blood (nM) | | | | | | |
| *Analyte* | *Control Average* | *Control Stdev* | *$d_7$-C12-BAC Average* | *$d_7$-C12-BAC Stdev* | *$d_7$-C16-BAC Average* | *$d_7$-C16-BAC Stdev* |
| $d_7$-COOH BAC C7 | 0.00 | 0.00 | 0.00 | 0.00 | 0.00 | 0.00 |
| $d_7$-COOH BAC C9 | 0.00 | 0.00 | 2.28 | 2.06 | 0.00 | 0.00 |
| $d_7$-COOH BAC C11 | 0.00 | 0.00 | 4.30 | 4.55 | 0.00 | 0.00 |
| $d_7$-COOH BAC C13 | 0.00 | 0.00 | 0.00 | 0.00 | 0.62 | 0.96 |
| $d_7$-COOH BAC C15 | 0.00 | 0.00 | 0.00 | 0.00 | 2.19 | 2.19 |

**Figure S1.** Odd Beta Oxidation Products in Male Blood and Liver

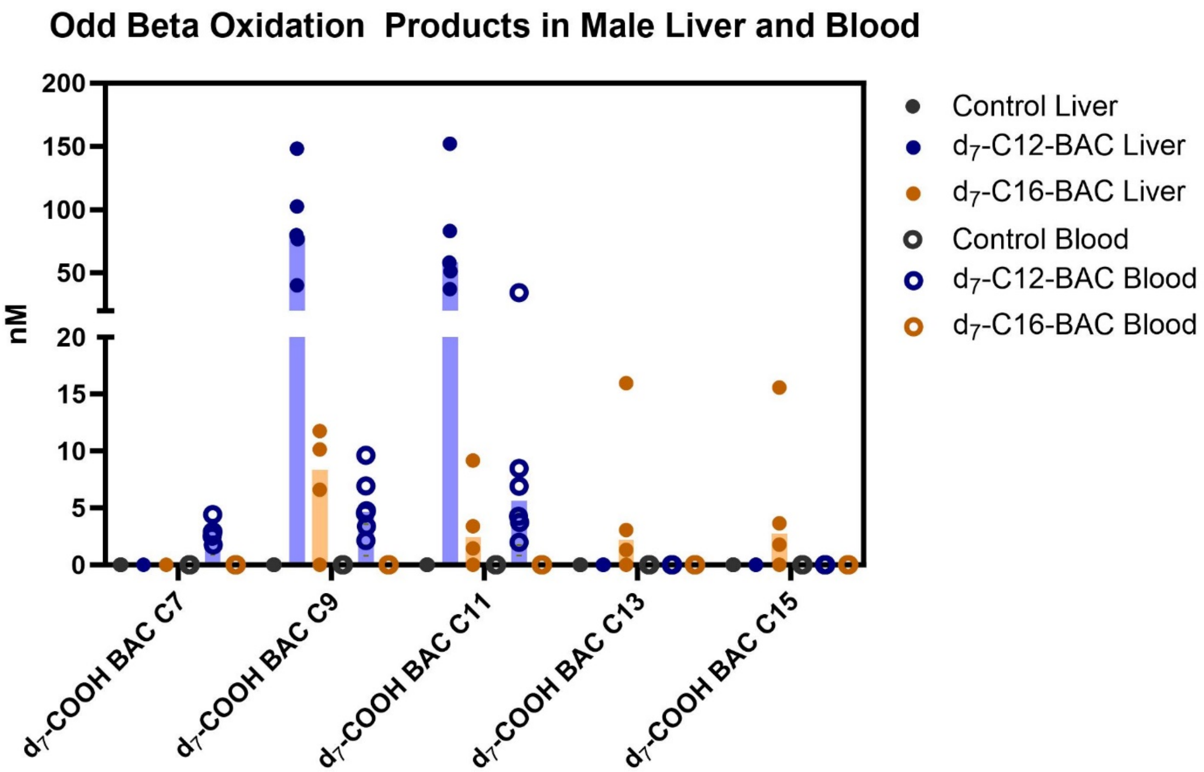

**Figure S2.** Odd Beta Oxidation Products in Female Blood and Liver

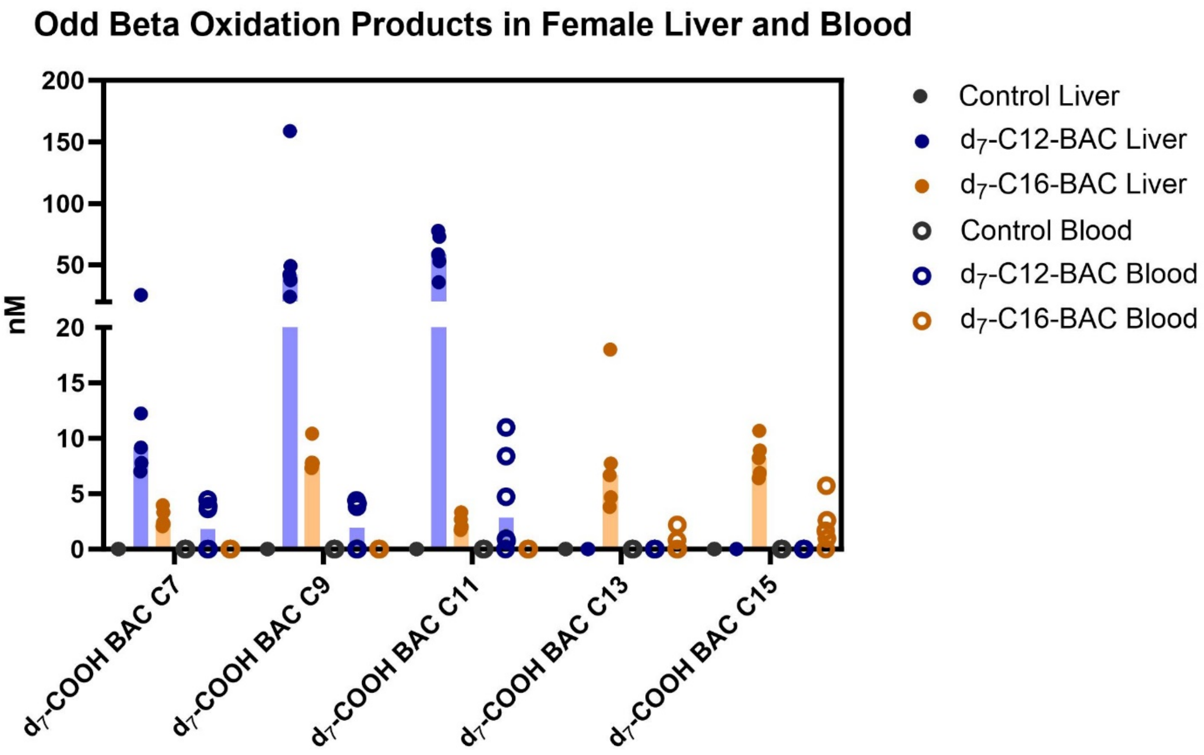

**Supplemental Table S7.** Average and Standard Deviations (Stdev) of odd chained beta-oxidation products in Male and Female Feces.

| Male Feces (µM) | | | | | | |
|---|---|---|---|---|---|---|
| *Analyte* | *Control Average* | *Control Stdev* | *$d_7$-C12-BAC Average* | *$d_7$-C12-BAC Stdev* | *$d_7$-C16-BAC Average* | *$d_7$-C16-BAC Stdev* |
| $d_7$-COOH BAC C7 | 0.00 | 0.00 | 1.48 | 1.12 | 0.66 | 0.33 |
| $d_7$-COOH BAC C9 | 0.00 | 0.00 | 12.53 | 9.42 | 2.72 | 1.27 |
| $d_7$-COOH BAC C11 | 0.00 | 0.00 | 113.17 | 51.22 | 4.83 | 2.11 |
| $d_7$-COOH BAC C13 | 0.00 | 0.00 | 0.00 | 0.00 | 22.52 | 8.36 |
| $d_7$-COOH BAC C15 | 0.00 | 0.00 | 0.00 | 0.00 | 95.72 | 31.53 |
| Female Feces (µM) | | | | | | |
| *Analyte* | *Control Average* | *Control Stdev* | *$d_7$-C12-BAC Average* | *$d_7$-C12-BAC Stdev* | *$d_7$-C16-BAC Average* | *$d_7$-C16-BAC Stdev* |
| $d_7$-COOH BAC C7 | 0.00 | 0.00 | 1.21 | 0.38 | 0.83 | 0.51 |
| $d_7$-COOH BAC C9 | 0.00 | 0.00 | 9.14 | 2.22 | 2.72 | 1.38 |
| $d_7$-COOH BAC C11 | 0.00 | 0.00 | 105.50 | 9.83 | 6.11 | 2.68 |
| $d_7$-COOH BAC C13 | 0.00 | 0.00 | 0.00 | 0.00 | 29.02 | 10.83 |
| $d_7$-COOH BAC C15 | 0.00 | 0.00 | 0.00 | 0.00 | 110.76 | 39.90 |

**Figure S3.** Odd Chained Beta Oxidation Products in Male Feces

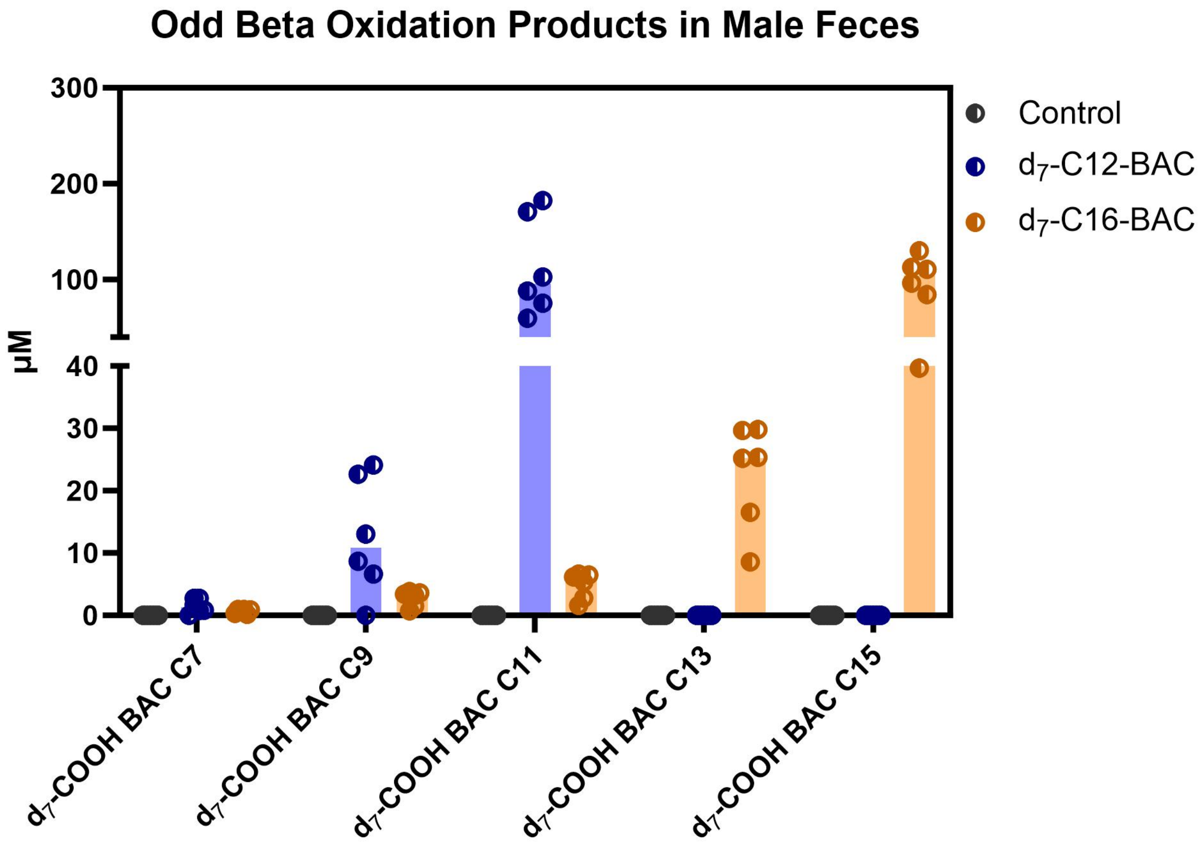

**Figure S4.** Odd Chained Beta Oxidation Products in Female Feces.

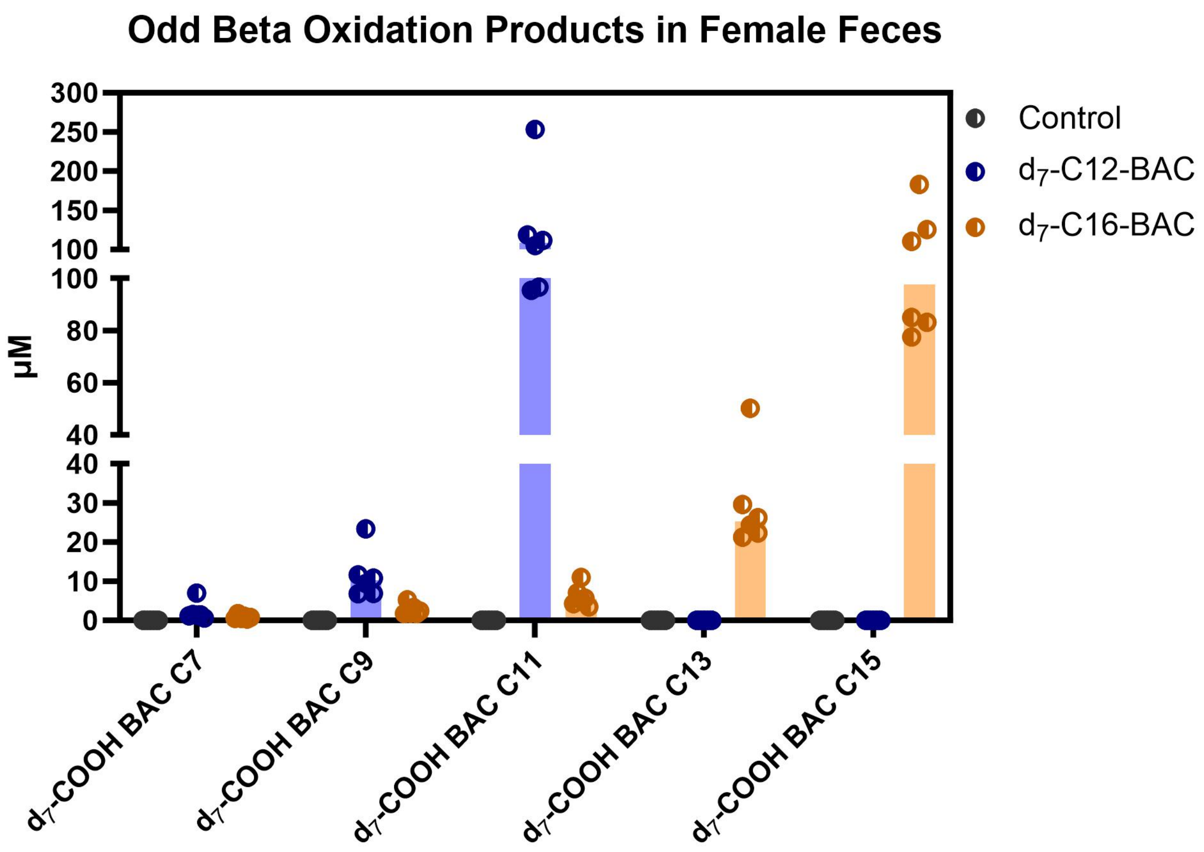

**Supplemental Table S8.** Precursor and Product Ion Transitions for BAC Quantitation

| Analyte | Retention Time (min) | Pre-cursor Ion m/z | Product Ion m/z |
|---|---|---|---|
| $d_7$-COOH C6 BAC | 1.5750 | 257.228 | 158.119/98.099 |
| $d_7$-COOH C7 BAC | 2.0950 | 271.16 | 172.133/98.099 |
| $d_7$-COOH C8 BAC | 2.6950 | 285.258 | 186.151/98.099 |
| $d_7$-COOH C9 BAC | 3.4950 | 299.275 | 200.17/98.099 |
| $d_7$-COOH C10 BAC | 4.2500 | 313.291 | 214.182/98.099 |
| $d_7$-COOH C11 BAC | 5.0050 | 327.31 | 228.201/98.099 |
| $d_7$-COOH C12 BAC | 5.6900 | 341.215 | 242.215/98.096 |
| (ω-1 OH)-$d_7$-C12-BAC | 5.7250 | 327.31 | 228.201/98.099 |
| (ω-OH)-$d_7$-C12-BAC | 5.9600 | 327.31 | 228.201/98.099 |
| $d_7$-COOH C13 BAC | 6.4450 | 355.34 | 256.23/98.099 |
| $d_7$-COOH C14 BAC | 7.1150 | 369.354 | 270.251/98.099 |
| $d_7$-COOH C15 BAC | 7.7800 | 383.37 | 284.261/98.099 |

| | | | |
|---|---|---|---|
| $d_7$-COOH C16 BAC | 8.4600 | 397.388 | 298.277/98.099 |
| (ω-1 OH)-$d_7$-C16-BAC | 8.7250 | 383.404 | 284.296/98.096 |
| (ω-OH)-$d_7$-C16-BAC | 9.0050 | 383.404 | 284.296/98.096 |
| $d_7$-C12-BAC | 10.1500 | 311.345 | 212.236/98.099 |
| $d_7$-C16-BAC | 13.4800 | 367.316 | 268.306/98.099 |
| COOH C8 | 2.7400 | 278.22 | 186.151/91.053 |
| COOH C10 | 4.2900 | 306.252 | 214.182/91.053 |
| COOH C12 | 5.7200 | 334.277 | 242.215/91.053 |
| 16 Da C12 | 5.8350 | 320.303 | 288.233/91.053 |
| COOH C14 | 7.1100 | 362.31 | 270.246/91.053 |
| COOH C16 | 8.4950 | 390.344 | 298.277/91.053 |
| 16 Da C16 | 8.8450 | 376.363 | 284.296/91.053 |
| C12 BAC | 10.1400 | 304.304 | 212.241/91.053 |

| | | | |
|---|---|---|---|
| C16 BAC | 13.5600 | 360.366 | 268.301/91.054 |
